# Dynamic Bayesian networks for integrating multi-omics time-series microbiome data

**DOI:** 10.1101/835124

**Authors:** Daniel Ruiz-Perez, Jose Lugo-Martinez, Natalia Bourguignon, Kalai Mathee, Betiana Lerner, Ziv Bar-Joseph, Giri Narasimhan

## Abstract

A key challenge in the analysis of longitudinal microbiome data is the inference of temporal interactions between microbial taxa, their genes, the metabolites they consume and produce, and host genes. To address these challenges we developed a computational pipeline, PALM, that first aligns multi-omics data and then uses dynamic Bayesian networks (DBNs) to reconstruct a unified model. Our approach overcomes differences in sampling and progression rates, utilizes a biologically-inspired multi-omic framework, reduces the large number of entities and parameters in the DBNs, and validates the learned network. Applying PALM to data collected from inflammatory bowel disease patients, we show that it accurately identifies known and novel interactions. Targeted experimental validations further support a number of the predicted novel metabolite-taxa interactions.

Source code and data will be freely available after publication under the MIT Open Source license agreement on our GitHub page.

**IMPORTANCE:** While a number of large consortia are collecting and profiling several different types of microbiome and genomic time series data, very few methods exist for joint modeling of multi-omics data sets. We developed a new computational pipeline, PALM, which uses Dynamic Bayesian Networks (DBNs) and is designed to integrate multi-omics data from longitudinal microbiome studies. When used to integrate sequence, expression, and metabolomics data from microbiome samples along with host expression data, the resulting models identify interactions between taxa, their genes and the metabolites they produce and consume, and their impact on host expression. We tested the models both by using them to predict future changes in microbiome levels, and by comparing the learned interactions to known interactions in the literature. Finally, we performed experimental validations for a few of the predicted interactions to demonstrate the ability of the method to identify novel relationships and their impact.

## INTRODUCTION

Microbiomes are communities of microbes inhabiting an environmental niche. The study of microbial communities offers a powerful approach for inferring their impact on the host environment, and their role in specific diseases and health. *Metagenomics* involves analyzing sequenced reads from the whole metagenome in a microbial community in order to determine a detailed profile of microbial taxa (1). More recently, additional types of biological data are being profiled in microbiome studies, including *metatranscriptomics*, which involves surveying the complete metatranscriptome of the microbial community (2), *metabolomics*, which involves profiling the entire set of small molecules (metabolites) present in the microbiome’s environmental niche (3), and *host transcriptomics*, which provides information about the levels of genes expressed in the host (4).

The goal of the second phase of the Human Microbiome Project (HMP) (5), called the integrative Human Microbiome Project (6), is to generate longitudinal multi-omics data sets as a means to study the dynamics of the microbiome and the host across select diseases, including preterm births, type 2 diabetes, and irritable bowel disorders.

A major challenge in microbiome data analysis is the integration of multi-omics data sets (7). Most multi-omic studies focus on a separate analysis of each omics data set without building a unified model (8). There have been some attempts (9, 10, 11, 12, 13) and tools to facilitate the analysis (14, 15), but there is still much room for improvement regarding reproducibility, flexibility, and biological validity (7, 16, 17).

Deep learning approaches for integrating multi-omics (18) have also been developed, but their lack of interpretability prevents these models from providing insights into the interplay of the different omics entities with the exception of MMvec (19), but it only combines metabolites and taxa. Even Partial Least Squares models have been used to facilitate this integration (20), but they have their own set of limitations depending on the underlying data generation model, and are prone to provide spurious results when applied to high-dimensional data (21).

In addition, microbiomes are inherently dynamic, and so to fully understand the complex interactions that take place within these communities, longitudinal microbiome data appears to be critical (22). Many attempts have been made to analyze data from longitudinal studies (23, 13, 12); however, these approaches do not attempt to study interactions between taxa. An alternative approach involves the use of dynamical systems such as the generalized Lotka-Volterra (gLV) models (24, 25), however the large set of parameters in these models diminishes their utility for probabilistic inference.

Previously, we have shown that probabilistic graphical models, specifically dynamic Bayesian networks (DBNs), can be used to study metagenomic sequence data from microbiome studies leading to models that can accurately predict future changes as well as identify interactions within the microbiome (26). However, these prior methods were only able to analyze a single omic data set. Here we present a new Pipeline for the Analysis of Longitudinal Multi-omics data (PALM), which, in addition to modeling metagenomics interactions can also incorporate time series metatranscriptomics, metabolomics and host expression data to learn an integrated model of microbiome-host interactions.

A number of challenges are associated with such large scale integration. First, modeling such data leads to a sizable increase in the size of the model and the number of parameters in the DBN, which grows as the product of the number of entities in each omics data set. More complex models makes the computation less tractable and harder to interpret. Additionally, such large number of nodes and parameters can lead to overfitting. PALM overcomes these challenge by restricting the set of allowable interactions (edges) between the omics entities based on sound biological assumptions and by relying on continuous representation and alignment to integrate a large set of observations when learning a specific model.

An additional challenge with modeling microbiomes is the difficulty of validating the model’s predictions. To address this, PALM uses *in silico* approaches employing multiple public databases (genomic sequence database and metabolic pathway database) and recently proposed software tools for the validation.

Applying PALM to Inflammatory Bowel Disease (IBD) data led to models that correctly predicted microbiome abundance levels and identified known and novel interactions. Statistical validations indicated that PALM can accurately recover known interactions and improved upon prior approaches. We also experimentally validated a few of the high scoring metabolite-taxa interactions predicted by the model.

## RESULTS

We developed a computational pipeline (PALM), presented in Figure 1, to process multi-omic microbiome data and infer their interactions. PALM first normalizes the data and then performs spline interpolation using continuous curves to enable imputation of missing time points and to overcome irregular sampling (Figure 1(b)). We next temporally align the data to correct for the different progression rates of each individual (Figure 1(c)) as well as filter out abnormal and noisy samples (Figure 1(d)). Alignment can be performed using either of the data types as we discuss below, and extrapolate the transformation to the other omics types. Our DBN learning algorithm utilizes prior knowledge to constraint the resulting model reducing overfitting and improving accuracy (Figure 1(e)). These dynamic constraints can be customized in the form of an adjacency matrix. Using the imputed, aligned data we learn a dynamic Bayesian networks (DBN) to model interactions within and between the different data types (Figure 1(f)). Finally, we validate the model predictive ability and the edges using a curated list of taxa-gene and taxa-metabolite interactions.

### Resulting Dynamic Bayesian network models

We used the Inflammatory Bowel Disease (IBD) cohort from the iHMP study (13) that followed 132 individuals over a year. These were profiled every two weeks on average, for different omics types. The preprocessing steps included filtering, interpolation, temporal alignment, variable selection, and removal of subjects with limited measured time points (see Methods for complete details). Based on these pre-processing steps, the resulting set used to learn the model consisted of 50 individuals across 101 microbial taxa, 72 genes, and 70 metabolites. In addition, for each host, the model includes 40 gene measurements profiled at two sites (ileum and rectum) from a single sampled time point), and the only environmental variable considered was the week in which the sample was obtained. We used this data set to learn multi-omic dynamic Bayesian models that provide information about interactions between taxa, genes and metabolites, and the impact of environmental variables and host transcriptomics on these entities over time. We used two sets of constraints; *Skeleton* and *Augmented* depicted in Supplemental File 1: Figure S1 and described in Section Constraining the DBN structure. The network with the complete IBD data set is presented in Supplemental File 2: Figure S2, with a total of two connections between the week of sample obtained and any other node, and Supplemental File 3: Figure S3, with no connections from the week variable, reinforcing the assumption that the system is in a steady state. For illustrative purposes of the capabilities of our methodology, we learned a DBN on a subset of the data set comprised solely of the 10 most abundant entities of each omic type as shown in Figure 2.

**FIG 1.**
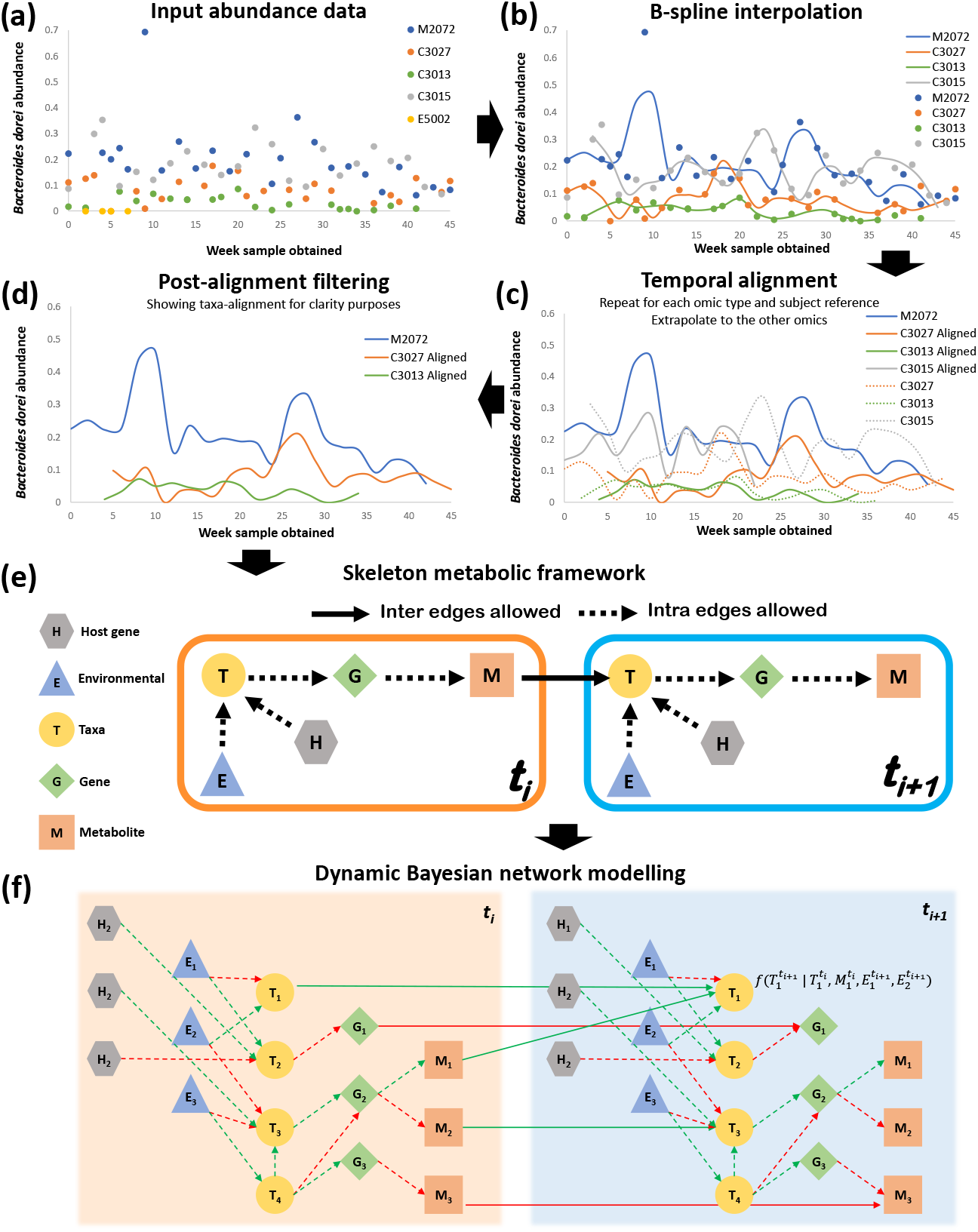
Computational pipeline proposed in this work. For simplicity, figure only shows microbial taxa *Bacteroides dorei* at each step in the pipeline from a set of five individual samples (subjects M2072, C3027, C3013, C3015, and E5002) of the IBD data set. (a) The relative abundance for each sample measured at potentially non-uniform intervals is the input of the pipeline. (b) Cubic B-spline curve for each individual sample. Subject E5002 (yellow) does not contain enough measured time points and was excluded from further analysis. The remaining smoothed curves enable principled estimation of unobserved time points and interpolation at specified intervals. (c) Temporal alignment of all taxa of each individual against the optimal reference subject (subject M2072 in blue). The learned warping function is extrapolated to all other omics data (e.g., genes and metabolites) of each subject. This process is then repeated, generating a different data set taking each omic as reference. (d) Post-alignment filtering of samples with a higher alignment error than a pre-defined threshold. Sample C3015 in grey was discarded. (e) Biologically-inspired *Skeleton* constraints imposed on learning the DBNs computed by PALM. The biological assumption is that at the current time (*t_i_*), the expression of host genes (hexagons) and the environmental conditions (triangles) affect the abundance of microbial taxa (circles), which impacts the expression of microbial genes (diamonds), which in turn dictates the metabolites (squares) released, and which finally impacts the abundance of taxa in the next time instant (*t*_*i*+1_). These restrictions are flexible and can be modified by the user as input to the pipeline. (f) Learning a two-stage DBN structure and parameters, where nodes correspond to either host genes, environmental variables, taxa, genes, or metabolites. Figure shows two consecutive DBN time slices *t_i_* and *t*_*i*+1_, where dotted lines connect nodes from the same time slice referred to as *intra edges*, and solid lines connect nodes between time slices referred to as *inter edges*. Biological relationships are annotated with the learned DBN parameters as either positive (green) or negative (red).

**FIG 2.**
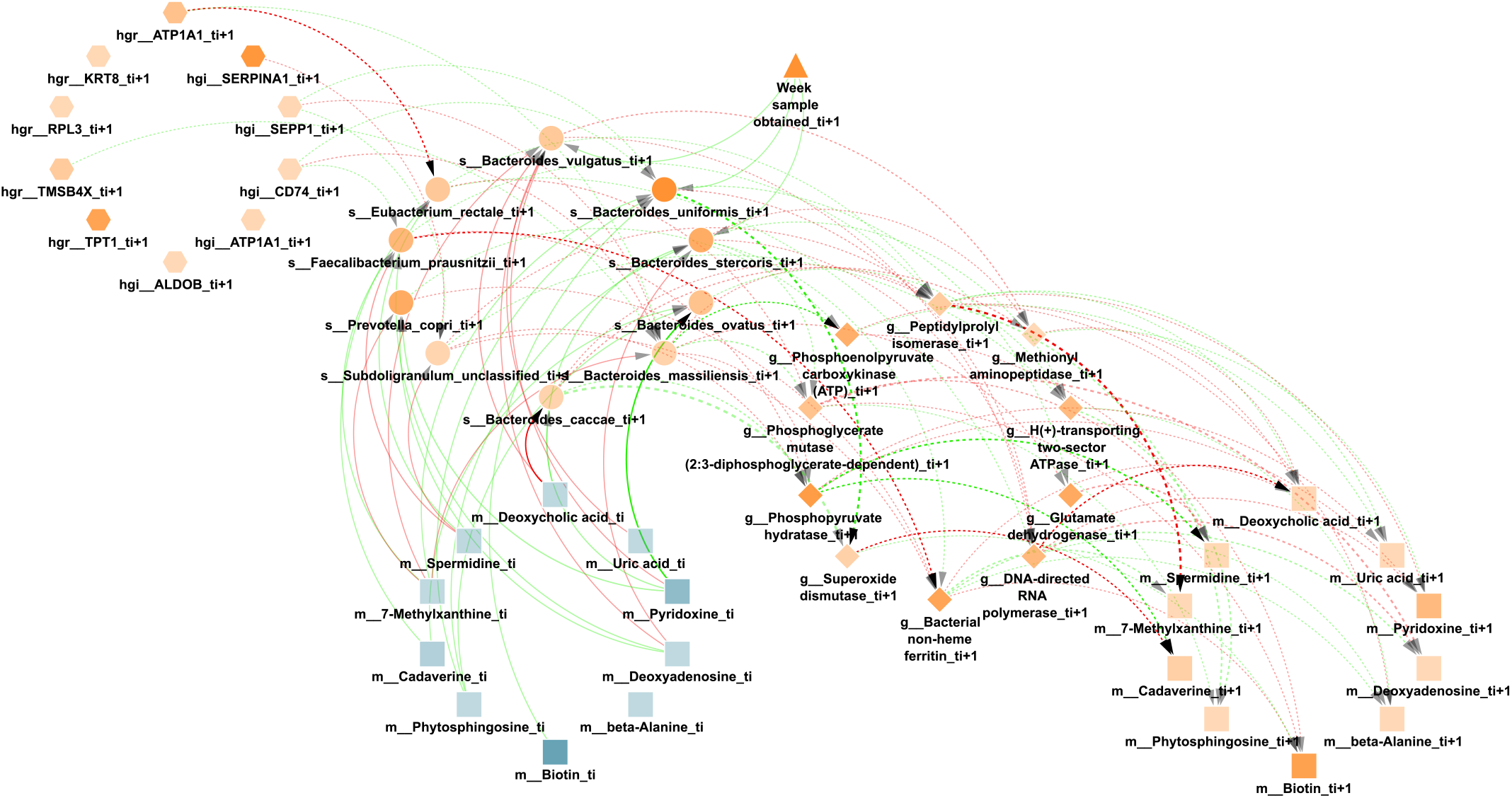
Learned DBN by PALM with the Skeleton constrains on the top 10 most abundant entities of each omic type and a maximum number of parents of 3. Nodes are either taxa (circles), genes (diamonds), metabolites (squares), host genes (hexagons), and environmental variables (triangles). The different node types have been grouped in different circles, their transparency is proportional to their average normalized abundance relative to that node type. While there are two consecutive time slices *t_i_* (blue) and *t*_*i*+1_ (orange), nodes with no neighbours and self loops were removed for simplicity. Dotted lines denote *intra edges* (i.e., directed links between nodes in same time slice), whereas solid lines denote *inter edges* (i.e., directed links between nodes in different time slices). Edge color indicates positive (green) or negative (red) temporal influence, and edge transparency indicates strength of bootstrap support. Edge thickness indicates statistical influence of regression coefficient after normalizing for parent values, as described in (26).

In the DBN figures, each node represents either a bacterial taxon, a gene, a metabolite, or an environmental variable; directed edges represent inferred temporal relationships between these nodes. On the supporting website we also provide a Cytoscape session with an interactive version of each network, together with the original files and a list of each edge learned weight for every network.

Supplemental File 2: Figure S2 shows the full network learned by PALM comprised of 284 nodes per time slice (101 microbial taxa, 72 genes, 70 metabolites, 40 host genes, and 1 environmental variable). To identify significant edges in the network we applied bootstrapping, which involves rerunning the method 100 times with each execution using a new data set created by randomly selecting, with replacement, as many subjects as there were in the data set. We next extracted all edges from all executions, resulting in 1077 distinct directed edges (470 inter edges and 607 intra edges). Among all the 1077 edges, we observed 362 (33%) negative interactions. Interestingly, a closer look at the learned DBN revealed that 79% (193 out of 243) of nodes with potential dependencies listed at least one negative interaction. Additionally, each edge is annotated with the percentage of bootstrap iterations in which it appears. Note that while there was considerable overlap between edges learned in each iteration, since we used the union of all the networks, the number of edges in the final network is larger than the number of possible edges for a single iteration (1077 vs. 284*3 = 852). While we mainly focus on the union since it leads to more novel predictions, analysis of the intersection leads to similar statistical results. The learned DBN with the Augmented framework is shown in Supplemental File 3: Figure S3.

### Evaluating the learned DBN model

We first performed a technical evaluation of the learned DBN model and compared it to models constructed by other existing methods (27, 26). The performance of each model was evaluated through leave-one-out cross-validation with the goal of predicting microbial composition using each learned model. Figure 3 represents the observed and predicted taxa composition for subject C3013.

Additionally, we explored the effects of several different temporal alignments using taxa, genes, or metabolites. In each iteration, the whole longitudinal microbial abundance profile of a single subject was selected as the test set, and the multi-omics data from all other subjects were used for building the network and learning model parameters. Next, starting from the second time point, we used the learned model to predict an abundance value for every taxon in the test set at each time point using the previous and current time points. Finally, we normalized the predicted values in order to represent the relative abundance of each taxon and measured the average predictive accuracy by computing the mean absolute error (MAE) for the selected taxon in the network. This process of predicting microbial composition was repeated for different combinations of multi-omics training data (including metagenomics, metatranscriptomics, metabolomics and host transcriptomics) on the aligned data sets, as well as unaligned data. A visual representation of the predicted trajectories for taxa- and gene-based alignment for subject C3028 is shown in Supplemental File 4: Figure S4. The average MAE for the taxa predictions of PALM on the IBD data set for a sampling rate of two weeks using a gene-based temporal alignment is summarized in Figure 4. Supplemental File 5: Figure S5 shows the average MAE of PALM across different alignments based on taxa, genes, and metabolites, respectively. Finally Supplemental File 11: Figure S10 and Supplemental File 12: Figure S11 show how MAE varies for different maximum number of parents for Augmented and Skeleton, respectively‥

**FIG 3.**
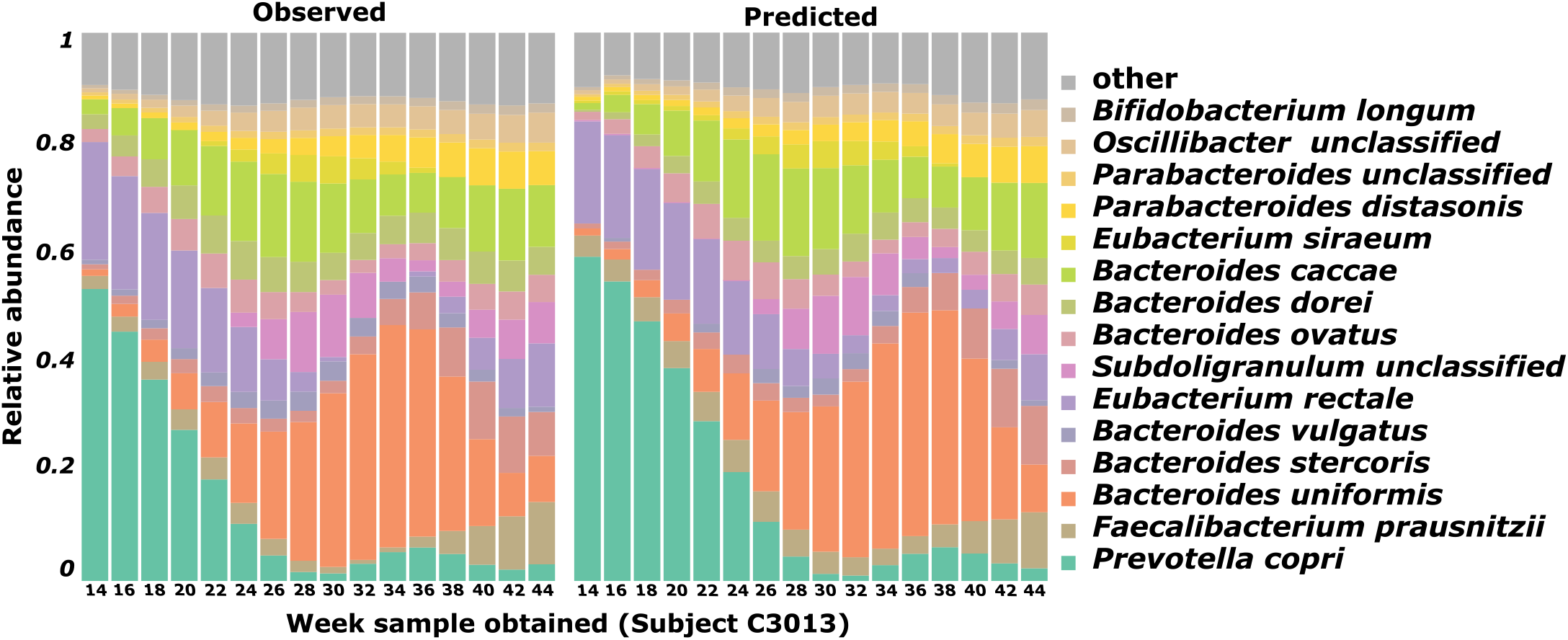
Comparison of observed versus predicted microbial composition trajectories. Figure shows the observed and predicted microbial composition trajectories for a representative aligned subject (C3013). Microbiota composition profile for this subject is comprised of the top 15 most abundant bacteria along with all remaining bacteria merged into the “other” category. The *y* axis corresponds to the relative abundance of each bacteria, while the *x* axis represents the original measured time point after alignment.

**FIG 4.**
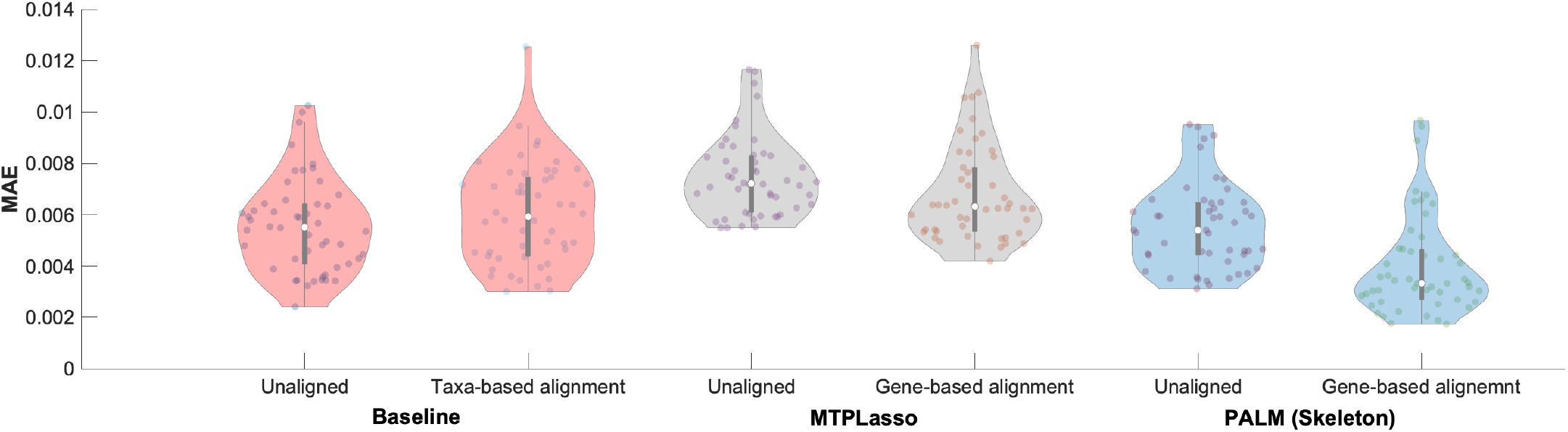
Comparison of average predictive accuracy between methods on the IBD data. Figure shows the MAE of our proposed DBN models against a baseline method using only metagenomic data and a previously published approach, MTPLasso, which models longitudinal multi-omics microbial data using a generalizded Lotka-Volterra (gLV) model for a sampling rate of two weeks which most closely resembles the originally measured time points. Figure also compares the performance of each method on the unaligned and aligned data sets.

We used this process to compare the multi-omics DBN strategy to the one that used only metagenomic data (26) referred to as Baseline on the unaligned and aligned IBD data, as well as MTPLasso (27) which models time-series multi-omics microbial data using a gLV model. In both cases, we used the default setup and parameters, as described in the original publications. As shown by Figure 4 our method outperforms Baseline and MTPLasso when using gene expression data for temporal alignment of microbiome samples. Specifically, when using gene expression data for alignment the MAE significantly dropped to 4.01E-03 when compared to a MAE of 6.03E-03 achieved using taxa alignment as indicated by a one-tailed unpaired t-test with null hypothesis that the means are equal and alternative hypothesis that population mean of method with gene expression-based alignment is less than mean of (baseline) taxa-based alignment method (p-value = 6.71E-07). Figure also shows that gene-based alignment significantly outperforms unaligned data regardless of the underlying method used. Similarly, in the case of taxa- or metabolite-based alignment, Supplemental File 5: Figure S5 shows that our method outperforms MTPLasso when all microbiome entities are used in the model (taxa: 5.93E-03 vs. 7.93E-03; metabolite: 5.82E-03 vs. 7.97E-03). Moreover, figure shows that our method outperforms Baseline (taxa: 5.93E-03 vs. 6.03E-03; gene: 4.01E-03 vs. 4.19E-03; metabolite: 5.82E-03 vs. 6.01E-03). Overall, our results suggest that gene expression data is more suitable for temporal alignment of multi-omics microbiome samples. This is consistent with previous findings which reported technical noise dominates the abundance variability for nearly half of the detected taxa in gut samples (28). Therefore, we have used gene-based alignment for the rest of the analysis discussed next.

In addition, since PALM can easily be extended to predict other omic types, we predicted the metabolite concentration and compared our results with MMvec (19) (Supplemental File 15: Figure S14). MMvec is the state of the art in metabolite prediction, and uses neural networks for estimating interactions between microbes and metabolites through their co-occurence probabilities. Our method significantly out-performs MMvec for all approaches tested (p-value = 4.10E-30 for a two-tailed paired t-test against the Augmented framework). This highlights the benefits of the time-series approach, which allows us to use the past and not only the present to predict future metabolite abundance profiles.

### Computationally validating predicted edges

We compiled a database of Taxon-Metabolite (*T* → *M*) and Taxon-Gene (*T* → *G*), and used that database to validate the predicted edges and score each model. A (*T* → *G*) interaction was added to the database if any strain of taxon *T* has gene *G* in its genome according to KEGG. For *T* → *M* we relied on the tool MIMOSA (29), that calculates the metabolic potential of each taxon for a particular data set. See Methods 4.7 for complete details

Each predicted interaction was either considered “validated” if it appears in the validation database, or “not validated” if it was not found, but the parent and child nodes were part of the database. Interactions predicted between taxa and/or metabolites not included in the database were not used in this analysis. We compared the results between the learned DBNs by PALM using the Skeleton and Augmented constraints, as well as a random network. To generate the random network, we used the same nodes in the multi-omic network and assigned the same number of edges as in the learned DBN by randomly selecting a parent and child from the possible interaction list (Supplemental File 1: Figure S1). This was repeated 1000 times, averaging the metrics over all random runs.

Figure 5(a) shows the validation comparison for edges of the form *T* → *G*. The Skeleton constraints were used to learn the networks. The learned DBN with the gene-aligned data set (green) was compared against a learned DBN with the data set that was not aligned (blue). As can be seen, the aligned data set results in networks that out-perform the networks from the unaligned data and random networks, with the precision difference increasing as the threshold increases. This indicates that the bootstrap score for an edge can serve as a way to determine its likely accuracy.

**FIG 5.**
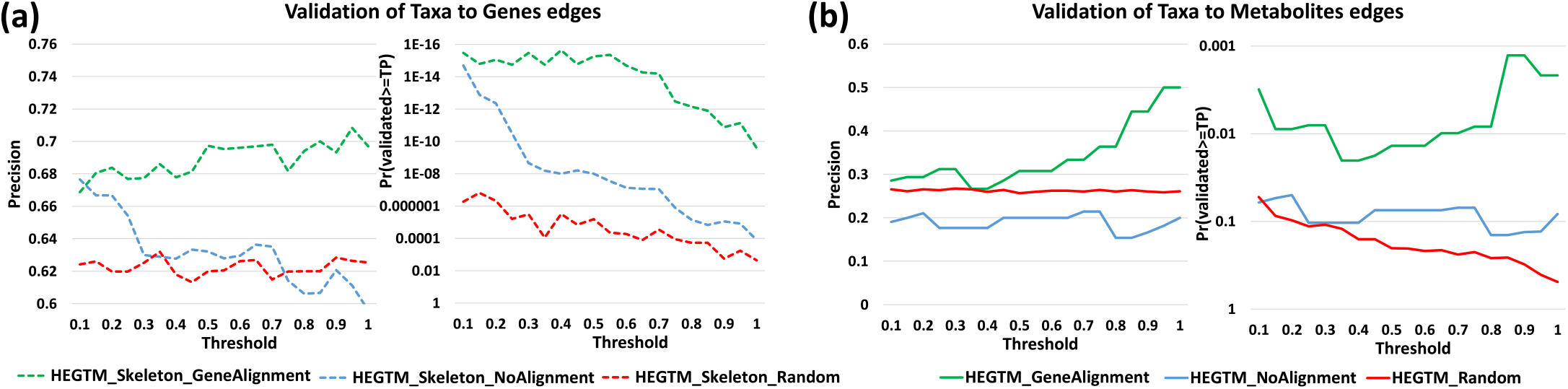
*In silico* validation results of the predictions of PALM for the IBD data set. The left part of each subfigure shows the precision (percentage of predicted edges that were validated) and the right part shows the probability of validating at least that many edges by chance (*y* axis in reverse logarithm scale so higher is better for both). The *x* represents the bootstrap value threshold that was used to select the edges included in the analysis. For example, for a threshold of 0.7, the score for edges that appear in more than 70% of the repetitions is shown‥ (a) Validation for *T* → *G* interactions (bacterial taxon expressing a gene) (b) Validation for *T* → *M* interactions (bacterial taxon consuming a metabolite)

Figure 5(b) shows the comparison for edges of the form *T* → *M*. For this, we can only use the network results from the Augmented constraints since no such edges are permitted when using Skeleton. Again, we observe better performance for the networks from aligned data when compared to the networks from unaligned data and random networks, with an improvement in performance for higher bootstrap thresholds. Note that for both *T* → *G* and *T* → *M*, the not aligned network does not even outperform the random network, highlighting the importance of the alignment step.

Supplemental File 6: Figure S6 shows the computational validation results of metabolites- and gene-based alignment, and for a varying maximum number of parents for both restriction frameworks. These results highlight our choice of alignment and maximum parents parameters. In addition, supplemental File 13: Figure S12 shows the validation of learned DBNs with gene-based alignments between sampling rates of 14d against 1d using the Skeleton framework. Finally, Supplemental File 14: Figure S13 shows the validation of the dataset normalized using *log-ratios* to circumvent bias in compositional data (30).

### Biological validation experiments

We performed experiments to validate a few of the interactions predicted by the DBNs. We focused on edges of the form *M* → *T*, i.e., edges where a metabolite is predicted to impact the abundance of a bacterial taxon. Such edges imply that the metabolite *M* promotes (or represses, depending on the sign) the growth of the bacterial taxon *T* under appropriate growth conditions.

We first sorted all predicted *M* → *T* interactions based on their confidence (product of normalized weight and bootstrap score). Next, we selected some of the top edges to validate taking into account the availability of the metabolites and taxa and the laboratory resources for growth experiments at our disposal. See Methods Section Laboratory validations of (metabolites to taxa) edges for details on this process. Based on these considerations we focused on two common model organisms, namely *Pseudomonas aeruginosa* and *Escherichia coli* and picked from the top predictions those involving any of these two taxa for validation. In addition, we learned independent densely sampled network (sampling rate of one day) to address the possible disconnect between the sampling rates of the DBN and the experiments. The results show that the learned networks are consistent when using the much denser sampling. Specifically, the skeleton gene-alignment networks shared 515 out of 676 (76%) of interactions with bootstrap score >= 0.5 (given the sparsity of the networks this is an extremely significant overlap). Only 10 bootstrap repetitions were used due to the computational resources needed. More importantly, all three positive controls tested in the wet lab were also found in the unaligned and aligned (metabolites-based) networks using Skeleton framework. Furthermore, two out of the three positive controls were also found in the learned networks from genes- and taxa-based alignments. Interestingly, none of the densely sampled networks learned the negative control interaction.

- 4-Methylcatechol (4-MC) → *Escherichia coli*
- 4-Hydroxyphenylacetate (4-HPA) → *Escherichia unclassified*
- D-Xylose → *Pseudomonas unclassified*

Standard lab strains *P. aeruginosa* PAO1 (31) and *E. coli* HB101 (32) were used in the laboratory experiments. The choice of chemicals used to verify was somewhat limited by commercial availability. A standard Luria Bertani (LB 20%) culture media was used to measure the bacterial growth curve (expressed as bacterial density, OD600) in the absence and presence of metabolites. Metabolites were added at the stationary phase when the bacteria were multiplying very slowly, mimicking a biofilm growth (33). As positive controls, the preferred carbon sources of *E. coli* and *P. aeruginosa*, glucose, and succinate, respectively, were chosen. Figure 6 shows the resulting growth curves of the microbes before and after adding the metabolites, and a control case without adding the metabolites (LB 20%). Confirming the predictions of our networks, D-xylose significantly enhanced *P. aeruginosa*, and 4-HPA and 4-MC significantly increased the *E. coli* growth. Regarding the controls, as expected D-xylose and glucose enhanced *E. coli*, and succinate enhanced *P. aeruginosa* whereas the negative control 1-Methylnicotinamide (1-MNA) did not.

**FIG 6.**
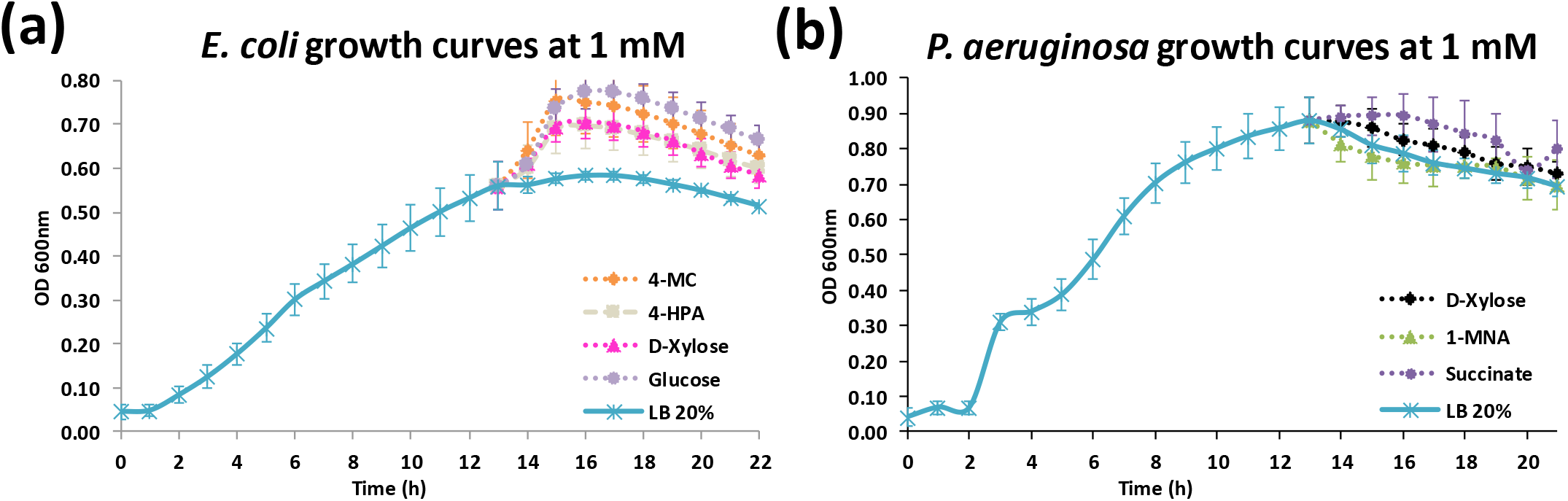
Growth curves at 1 mM. In this figure different metabolites were introduced at 1mM concentration at the end of the exponential phase (0-14h). Figure shows the growth curves after all data points were averaged over 10 replicates. (a) *E. coli*, with glucose and D-xylose as positive controls. (b) *P. aeruginosa*, with succinate as positive control and 1-MNA as negative control.

The p-values for all observations can be seen in Table 1, where a two-tailed paired t-test was executed for the three time points with the highest difference from the baseline. For more details on the experimental settings please refer to Methods Section Laboratory validations of (metabolites to taxa) edges, including Supplemental File 7: Figure S7 and Supplemental File 8: Table S1 for growth results at a lower concentration of 0.2 mM.

**Table 1.** Effect of 1 mM metabolites on bacterial cell density. Taxa density appears in black (OD600), while the p-values are inside parenthesis. Red p-values represent a signifficant difference compare to LB 20% (p<0.05). Green p-values represent a non-signifficant difference from LB 20%.

| Escherichia coli HB101 |  |  |  |
| --- | --- | --- | --- |
| Met. | 16h | 17h | 18h |
| LB | 0.583 | 0.584 | 0.576 |
| 4-MC | 0.750 (0.0002) | 0.741 (0.0003) | 0.724 (0.0003) |
| 4-HPA | 0.697 (0.0005) | 0.692 (0.0005) | 0.677 (0.0004) |
| D-xylose | 0.704 (0.0017) | 0.699 (0.0016) | 0.684 (0.0019) |
| Glucose | 0.774 (0.0005) | 0.773 (0.0003) | 0.758 (0.0003) |

**TABLE 1** Effect of 1 mM metabolites on bacterial cell density. Taxa density appears in black (OD600), while the p-values are inside parenthesis. Red p-values represent a significant difference compare to LB 20% ( $p < 0.05$ ). Green p-values represent a non-significant difference from LB 20%.
| Pseudomonas aeruginosa PAO1 |  |  |  |
| --- | --- | --- | --- |
| Met. | 15h | 16h | 17h |
| LB | 0.811 | 0.787 | 0.760 |
| D-Xylose | 0.860 (0.0189) | 0.825 (0.0395) | 0.810 (0.0317) |
| 1-MNA | 0.782 (0.0717) | 0.761 (0.1551) | 0.754 (0.7547) |
| Succinate | 0.893 (0.0716) | 0.893 (0.1195) | 0.870 (0.0172) |

## DISCUSSION

Previous microbiome studies focused primarily on metagenomics sequence data. More recent data sets are much richer, notably including host and bacterial gene expression, and metabolomics data. The ability to integrate these multi-omics, longitudinal data remains a major challenge for microbiome analysis.

Here we have presented PALM, a new approach based on a temporal normalization using continuous curve alignment followed by DBN modeling. Our method first represents each time series using continuous curves and then aligns them using a reference time series. Next, we sample the aligned curves uniformly and learn a DBN model that combines data from taxa, host genes, bacterial genes, and metabolites. Edges in the DBN represent predicted interactions between the entities and can be used to explain changes in the microbiome over time.

Applying our methods to data from IBD patients, we show that multi-omics DBNs can successfully predict taxa abundance at future time points, thus improving on models that do not use all available data and on previous methods developed for modeling temporal taxa interactions. We curated validations for taxa-to-metabolite and taxa-to-gene interactions; edges predicted by the learned DBNs significantly intersect these interactions. Finally, we experimentally tested and validated select predictions of metabolite → taxa relationships. We have ignored the case-control structure of the IBD dataset for this work, but the framework can be easily used for that purpose either by learning different models for each disorder or by adding a ‘diagnosis’ node to the network and studying its outgoing edges.

Microbiome interaction databases are critical for evaluating learned DBNs, but appear to be incomplete. More complete databases of validated interactions would help validate computational methods for this task. The laboratory validations show a viable way to validate some of the interactions. However, they could also be improved by attempting to recreate more realistic conditions for the experiments and could be enhanced to validate other omics observations as well. Our models suggest that certain metabolites can be used as predictors of the abundance of taxa. Our interpretation is that bacteria are consuming these metabolites, which causes fluctuations in their abundance. But other indirect effects could also be possible

Comparing DBNs constructed using different omics data allows for an important kind of inference (Supplemental File 9: Figure S8). According to this, in the DBN built using only metagenomics data, the edge *Streptococcus parasanguinis* → *Pseudomonas unclassified* appears with a high confidence (bootstrap score of 1). In the multi-omic DBN the following chain of interactions can be found: *Streptococcus parasanguinis* (T) → rna polymerase (G) → D-Xylose (M) → *Pseudomonas unclassified* (T). It is important to note that though DBN edges may not imply causal relationships, the *in silico* validation process described in this paper supports the above relationships. Finally D-Xylose → *Pseudomonas unclassified* was validated experimentally (Section Biological validation experiments). Thus, comparing DBNs before and after adding additional multi-omics data can “unroll” and “explain” relationships between taxa.

Our alignment and DBN methods are implemented in Python and Matlab correspondingly. The source code and data set used can be obtained from the link on the cover page to reproduce the findings of this paper, together with the networks learned and interactions predicted, sorted by relevance.

## MATERIALS AND METHODS

Below we describe the computational pipeline, PALM, developed to integrate and model the interactions between the different types of omics.

### Data

To test PALM’s proposed analysis pipeline, which combines temporal alignment with Bayesian network learning and inference for multi-omics microbiome data, we used the Inflammatory Bowel Disease (IBD) cohort from a study that included 132 individuals across five clinical centers (13). During a period of one year, each subject was profiled (biopsies, blood draws, and stool samples) every two weeks on average. This yielded temporal profiles for metagenomes, metatranscriptomes, proteomes, metabolomes and viromes across all subjects. Although the metatranscriptomics data summarized functional profiling via HUMAnN2 (34) at species-specific and species-agnostic quantification of gene families, EC enzyme modules, and pathways, we solely focus on the metabolic enzymes (i.e., EC enzyme modules) quantification at the community level. Additionally, for each subject, host- and microbe-targeted human RNA sequencing was yielded from biopsies collected at initial screening colonoscopy sampled at three locations (colon, ileum and rectum); however, only ileum and rectum data was used in this study. All data sources are fully described and available at https://ibdmdb.org/.

Each data set was associated with the week when it was sampled, except for the host transcriptomics where only a single biopsy was obtained at each location (ileum and rectum). Therefore, host transcriptomics data were used as static variables in the DBN.

### Data pre-processing

First, for each subject, the different omic profiles (taxon, gene, and metabolite) were normalized separately such that each omic type sum up to 1. Next metabolites and genes were scaled to match the mean of the taxa solely for visualization purposes. We note that the learned DBNs with and without scaling were exactly the same (data not shown). In order to account for the common pitfalls when comparing relative abundance across samples as highlighted by recent studies (35, 30, 36), we also separately normalized microbial taxa data using *log-ratios* which have been shown to circumvent bias in compositional data (30). Specifically, we used *Faecalibacterium prausnitzii* as the reference species. Next, for each individual with *n* > 1 different longitudinal samples *s*_1_,…, *s_n_*, we computed the log-ratio as

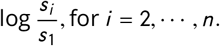

Additionally, RNA-seq data from ileum and rectum of each host was analyzed using DESeq2 (37) and TPM values. As a proof-of-concept, for each body site, we selected the top 20 genes with the highest variance across all subjects. We note that this set of genes is limited as previous studies have reported over 1000 differently expressed genes for IBD individuals at these locations when compared to individuals without IBD (13). In the case of metabolomics data, metabolites without an HMDB correspondence were removed. Then, we filtered out metabolites for which the mean intensity was less than 0.1%, or had zero variance over the originally sampled time points. Next, we performed temporal alignment of time series data from individuals as described in Lugo-Martinez *et al.* (26). For this, we need to represent each discrete time series using a continuous function. Here we used B-splines for fitting continuous curves to the time-series multi-omic data profiled from each subject, including the microbial composition, gene expression, and metabolic abundance. To improve the accuracy of the reconstructed profiles, we removed any sample that had less than five measured time points in any of the multi-omics measurements. Although the IBD data is composed of 107 individuals with taxonomic profiles, 77 individuals with gene expression profiles and 80 individuals with metabolic profiles, where each omic profile has at least 5 longitudinal measurements, there are exactly 62 individuals whose trajectories have at least 5 measured timepoints for all three omic types. This set is further reduced to 51 individuals when cross-referenced with the gene expression profiles derived from host transcriptomics data. Therefore, the final set is comprised of 51 individual multiomic time series used for further analysis.

### Temporal alignments

Given longitudinal samples from different subjects, we cannot expect that the rates at which various multi-omics levels change would be exactly the same between these individuals (38). To facilitate the analysis of such longitudinal data across subjects, we first align the time series from the microbiome samples using the microbial composition profiles. As described earlier, these alignments use a linear time transformation function to warp one time series into a common, representative sample time series used as the reference (26). While prior alignment methods relied on taxa information, when multi-omics data is available, PALM can use other genomic information for the alignment. Specifically, here we also tested the use of gene expression and metabolite abundance profiles for determining accurate alignments of patients. As we show, by using a better omics data type the resulting DBNs can more accurately capture and predict taxa-metabolite and taxa-gene relationships.

For each omics data (i.e., taxa, genes, or metabolites) we select an optimal reference sample from the 51 time series as follows: we generated all possible pairwise alignments between them and selected the time series that resulted in the least total overall error in the alignments. We then searched for abnormal and noisy samples from the resulting set of alignments as follows: (1) computed the mean *μ* and standard deviation *δ* of the alignment error, and (2) removed all samples from an individual where alignment error exceeded *μ* + (2 × *δ*), as previously described in Lugo-Martinez *et al.* (26). However, we did not remove any of the 51 time series samples as none of the aligned profiles displayed an alignment error satisfying these constraints across all three omic types. Figure 1(a)-(d) shows the overall alignment process of *Bacteroides dorei*, from the taxa-based alignment perspective.

Given an individual’s warped/aligned time series over a specific omic type, the other multi-omics data were incorporated as follows: the same transformation applied to the aligned sample was applied to all the complementary multi-omics time series data. The resulting set used for the modeling comprised of 50 individual-wise heterogeneous alignments involving 101 microbial taxa, 72 genes, and 70 metabolites. This smaller number of attributes was used because learning a Bayesian network is NP-Hard (39, 40) and has an exponential run-time with the number of features, however there is no imposed restriction on the number of features.

### Dynamic Bayesian network models

Using the aligned time series multi-omics data, we next learned graphical models that provide information about the relationships between the different omics (taxa, genes, metabolites, host-genes) and environmental (exogenous) variables. In PALM, we extend the DBN model proposed in Lugo-Martinez *et al.* (26) to account for multi-omics microbiome data with the goal of inferring the temporal relationships between the heterogeneous entities in a microbial community. A DBN is a directed acyclic graph where, at each time slice, nodes correspond to random variables of interest (e.g., taxa abundance, gene expression, age, etc.), and directed edges correspond to their conditional dependencies in the graph. These edges are defined as either: *intra edges* connecting nodes from the same time slice, or *inter edges* connecting nodes between consecutive time slices. In our DBN model, only two slices are modeled and learned, as shown in Figure 1(e).

In PALM, our DBN models encode five types of nodes: (i) taxon abundance, (ii) gene expression, (iii) metabolite concentration, (iv) host gene expression, and (v) sample metadata information. The first three types represent continuous variables, whereas the last two types can be either discrete or continuous. For our DBNs, we use the formalism of conditional Gaussian Bayesian networks (41) to take advantage of its ability to seamlessly integrate discrete and continuous variables in a single probabilistic framework. Formally, let Θ denote the set of parameters for the DBN and *G* denote a specific network structure over discrete variables (denoted as ∆) and continuous variables (denoted as Ψ) in the multi-omics microbiome study. The joint distribution *P* (∆, Ψ) can be decomposed as

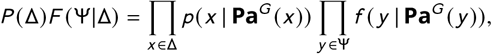

where *P* denotes a set of conditional probability distributions over discrete variables, *F* denotes a set of linear Gaussian conditional densities over continuous variables, and **Pa**^*G*^ (*X*) denotes the set of parents for variable *X* in *G* (42, 26). In particular, continuous variables are modeled using a Gaussian with the mean set based on a regression model over the set of continuous parents as follows

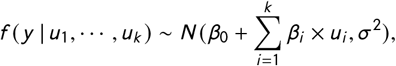

where *u*_1_,…, *u_k_* are continuous parents of *y*; *β*_0_ is the intercept; *β*_1_,…, *β_k_* are the corresponding regression coefficients for *u*_1_,…, *u_k_*; and *σ*^2^ is the standard deviation. In the case that *y* has discrete parents then we need to compute coefficients 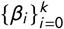 and standard deviation *σ*^2^ for each discrete configuration. This Gaussian regression model is appropriate for modeling errors, and is at least partially a way to deal with both normalization impact and measurement noise. As highlighted in Figure 1(e), the conditional linear Gaussian density function for variable 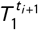 denoted as 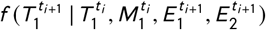, is modeled by

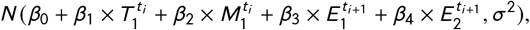

where Θ = {*β*_1_, *β*_2_, *β*_3_, *σ*^2^} are the DBN model parameters. Here, we infer the parameters Θ by maximizing the likelihood of the longitudinal multi-omics data *D* given our regression model and known structure *G*.

The problem of learning the DBN is expressed as finding the optimal structure and parameters

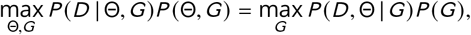

where *P* (*D* | Θ, *G*) is the likelihood of the data given the model. Since the likelihood of a structure increases as the number of edges increases, one must effectively find the structure that maximizes the likelihood of the data while penalizing overly complex structures. As in Lugo-Martinez *et al.* (26), we maximize *P* (*D*, Θ | *G*) for a given structure *G* using maximum log-likelihood estimation (MLE) combined with BIC score defined as

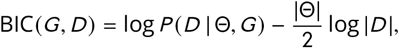

where |Θ| is the number of DBN model parameters in structure *G*, and |*D* | is the number of observations in *D*. This approach enables an effective way to search over the set of all possible DBN structures while favoring simpler structures. Furthermore, this approach has been shown to outperform Bayesian-Dirichlet scores, which require prior knowledge and can be sensitive to parameters and improper prior distributions (43, 44, 26).

### Constraining the DBN structure

An important innovation in PALM lies in the structure constraining of the network to conform to our proposed metabolic framework that ensures the desired flow of interactions. These constraints (in the form of a matrix received as an input to the function) only allow edges between certain types of nodes, highly reducing the complexity of searching over possible structures and preventing over-fitting. Note that these constraints can be easily modified by the user such as adding more data types, or different restrictions in the input file containing the adjacency matrix. Specifically, we allowed intra edges from environmental and host transcriptomics variables to microbial taxa (abundance) nodes, from taxa nodes to gene (expression) nodes and from gene nodes to metabolites (concentration) nodes. All other interactions within a time point (for example, direct gene to taxa) were disallowed. We also allowed inter edges from metabolites to taxa nodes in the next time point, and *self-loops* from any node 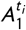 to 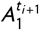, except for environmental or host transcriptomics variables for which no incoming edges were allowed (host genes were only measured at a single time point so no incoming temporal edges were allowed for them). These restrictions referred to as the *Skeleton* and depicted in Supplemental File 1: Figure S1(a) reflect our understanding of the basic ways the different entities interact with each other, i.e., environmental and host gene expression variables are independent variables, taxa express genes, which are involved in metabolic pathways; finally, the metabolites impact the growth of taxa (in the next time slice).

We also learned DBNs using a less constrained framework referred to as *Augmented* as shown in Supplemental File 1: Figure S1(b). Unlike Skeleton, the Augmented framework also allowed direct edges between taxa and metabolites to account for cases where noise or other issues related to profiling of genes can limit our ability to indirectly connect taxa and the metabolites they produce. Supplemental File 1: Figure S1 summarizes each framework in the form of an adjacency matrix. Note that other constraints such as requiring that taxa could only connect to genes present in their genome were not imposed since genomics reference databases are not always complete and so they may lead to missing key interactions.

We used a greedy hill-climbing approach for structure learning where the search is initialized with a network that connects each node of interest at the previous time point to the corresponding node at the following time point. Next, nodes are added as parents of a specific node via intra or inter edges depending on which valid edge leads to the largest increase of the log-likelihood function beyond the global penalty incurred by adding the parameters as measured by the BIC score approximation.

Every network was bootstrapped by randomly selecting with replacement as many subjects as in the data set, and learning a different network 100 times. Although we explore multiple values as the maximum number of possible parents for each node (see Supplemental Figures S6, S9, S10, and S11 for results with different maximum number of parents), unless otherwise stated, the maximum number of possible parents was fixed to 3. The networks were then combined, and the regression coefficient of the edges was averaged. Each edge was also labeled with the bootstrap support (percentage of times that edge appears). Each repetition was set to run independently on a separate processor using Matlab’s Parallel Computing Toolbox. Other parallel implementations include parallelizing the cross-validation computation of the inference error and each independent alignment error calculation using Python’s Parallel library.

### Validating DBNs

A major challenge in building models of biological interactions lies in developing methods to validate them and in providing confidence measures. Since DBNs are generative models, one approach is to predict time series using previous time points and thus to achieve cross validation (26). Such technical validations, while informative, could be thought as of “black-box” validation, and do not shed light on the accuracy of specific edges and interactions predicted by the model that we are interested in.

We broadly discuss approaches to validate the types of edges present in the DBN (see Figure 1(e)), which are the parameters learned by the model, and hence closer to “white-box” validation. Edges from taxa to genes can be circumstantially validated by verifying that (a) the taxon presence is guaranteed by its non-zero abundance, (b) the taxon genome has the gene, and (c) the gene is expressed. PALM, therefore, handles this using the *in silico* validation strategies mentioned below in Section 4.7. Similarly, edges from genes to metabolites or taxa to metabolites could potentially be validated.

The challenge is in validating edges from metabolites to taxa, for which an *in silico* approach is unlikely to work since no such database has been compiled to the best of our knowledge. In Section 4.8, we propose a validation approach involving laboratory experiments.

### *In silico* validation of DBN edges

*In silico* validations of DBN edges are handled by verifying the information against a database of known interactions between taxa to genes and/or taxa to metabolites. Unfortunately, no such comprehensive database exists. For example, highly curated databases such as HMDB (45), MetaCyc (46), or the findings of the large scale study of Maier et al. (2019) (47) turned out to be inadequate since the intersection of their contents with the species and metabolites in our networks was too small.

To assist in the validation of taxa-metabolite (*T* → *M*) edges in our networks, we relied on the tool MIMOSA (29). MIMOSA calculates the metabolic potential of each species, i.e., the capability of a species to produce a metabolite under the conditions of the data set. The list of all taxon-metabolite pairs from our DBNs that resulted in a positive score in MIMOSA was used as a validation database.

For taxa-gene (*T* → *G*) validations, we used KEGG to build a validation database of bacterial taxa and the genes present in their genomes. To keep this database small, we only used taxa and genes present in our network. If multiple strains were available for a bacterial species, then all genes from each strain were aggregated. The one-time creation of a local validation database also speeded up our computations considerably.

To calculate the statistical significance of validated interactions compared to a null model, a Poisson-Binomial distribution test was executed. The main reason that a simple binomial test cannot be performed is the differences in the in-degree distribution between different nodes in the validation set (essential metabolites or genes would have a high probability of being connected to any given bacteria in the validation database). Because some nodes have many more validated interactions when compared to others, a uniform model for each edge does not accurately capture the null probability of selecting such an edge. This was solved with the function ppoisbinom from the R package poisbinom (48), which gives the cumulative distribution function of the probability of validating by chance at least as many interactions as the number of true positives, where each possible interaction has a different probability of being selected. The validation precision of the network was also calculated as the percentage of validated interactions from the ones predicted, even though this homogeneous metric ignores the differential significance of each interaction. We did not calculate the validation recall because in many cases a false positive cannot be distinguished from a true interaction that was simply not recorded in the database, and could lead to misleading scores.

### Laboratory validations of (metabolites to taxa) edges

Wet lab experiments were carried out to validate predicted *M* → *T* interactions. Testing each such edge is not a feasible proposition. We first sorted all predicted *M* → *T* interaction based on their confidence, which we defined as the value of |*normalize* (*weight*)| * *bootstr ap*. We applied this operation to the three parents Skeleton for the gene-aligned and noalignment networks. The normalization was performed to counteract the differences of the abundance between the parent and child nodes following (26). We narrowed it down to edges that involved the species *P. aeruginosa* or *E. coli* because of the ready availability of these species and the expertise and facilities available to us in our laboratories. Then, we combined and sorted all interactions from gene-alignment and no-alignment and selected the top interaction involving *P. aeruginosa* and the top two interactions involving *E. coli* to validate. The full list of interactions along with their confidence scores can be seen in the Networks folder of the supplementary material. For positive controls we selected metabolites known to enhance growth, and as negative control we selected one metabolite that was not connected to the taxon in any of our learned networks.

The goal of the experiments was to validate a *M* → *T* edge by studying the impact of the metabolite *M* on the growth of taxon *T*. While the experimental set up does not recreate the conditions of the interaction in the microbiome, we consider this an important step in the right direction. As with the *in silico* validations, the laboratory validation confirms that the inferred interaction is a strong possibility. We selected three predicted interactions involving readily available bacteria and metabolites from the generated networks. The experiments were performed by growing relevant taxa in isolation, and adding the relevant metabolite to measure impact on growth. These metabolites were expected to positively impact the growth of the taxon because of the edge between metabolite concentration and taxon abundance. To address the apparent disconnect of the time scales between the dataset (two weeks) and the laboratory experiments (several hours), we also subsampled the original dataset with a sampling rate of one day and learned independent networks, making sure the tested interactions were also in these densely sampled networks.

After plotting the growth curves with the bacterium and metabolite in question, we assessed if each metabolite was enhancing/inhibiting the taxon growth using a two-tailed paired t-test when compared to growth without the metabolite.

#### Preliminary experiments

Three preliminary experiments were run that paved the way for the final experiment. The bacterial strains used *Escherichia coli* HB101 (32) and *Pseudomonas aeruginosa* PAO1 (31) were routinely cultured in Luria Bertani (LB 20%) broth (5 g tryptone, 10 g sodium chloride, and 5 g yeast extract per liter) or agar (LB broth with 1.5% agar) (Difco, NJ, USA). Growth curve assays were performed in media supplemented with the metabolites at 37°C. For the three preliminary experiments, we attempted to closely mimic limited nutrient environment.

1. Experiment 1, tested 0.2 mM:

- Minimal Media (MM; gL−1: (NH_4_)_2_SO_4_, 2.0; K_2_HPO_4_, 0.5; MgSO_4_ · 7H_2_O, 0.2; FeSO_4_ · 7H_2_O, 0.01, pH 7.2±0.2).
- MM + Glucose
2. Experiment 2: LB 20%, tested 0.2, 1.0 and 2 mM
3. Experiment 3: LB 20%, tested 0.2, and 1.0 mM *E. coli* was grown in the presence of 4-methylcatechol (4-MC, C_7_H_8_O_2_) and 4-hydroxyphenylacetate (4-HPA, C_8_H_8_O_3_). *P. aeruginosa* was grown in the presence of D-xylose (C_5_H_10_O_5_), and 1-methylnicotinamide (1-MNA, C_7_H_9_N_2_O_+_).

1. Experiment 1 (0.2 mM of metabolites)

- The cells reached stationary phase at a very low OD; suggesting that this is not the right media to be used.
- No effect on the exponential phase.
- Any effect of the compound seen at the stationary phase.
2. Experiment 2 (0.2, 1 and 2 mM of metabolites)

- The cells reached stationary phase at a higher OD.
- No effect on the exponential phase.
- 2 mM is lethal
3. Experiment 3 (0.2, and 1 mM of metabolites)

- The cells reached stationary phase at a higher OD.
- No effect on the exponential phase.
- Effect is seen during the stationary phase

#### Final experiment

Because no significant difference was observed in the exponential growth rate and consequently the doubling time in all the conditions tested for both *E. coli* and *P. aeruginosa*, a new test was run in which the metabolites were added at the beginning of the stationary phase to test its effect on it.

The growth of *E. coli* was monitored hourly in the absence (control) and presence of 4-MC, 4-HPA, D-Xylose and glucose at 0.2 and 1 mM. Glucose and D-Xylose were used as enhancer positive controls. At the lower concentration (0.2 mM) compared to the control (LB 20%), 4-HPA has no effect and 4-MC, D-Xylose and glucose are enhancing (Supplemental File 7: Figure S7(a)). The compound 4-HPA has no effect at low concentration, however at 1 mM, there is a significant enhancing effect starting at early stationary phase. At the highest concentration all metabolites produce an enhancer effect statistically significant (t-test, p<0.05), there is also a more pronounced enhancer effect of 4-MC and glucose compared to D-Xylose and 4-HPA (Figure 6).

The growth of *P. aeruginosa* PAO1 was monitored hourly in the absence (control) and presence of D-xylose, 1-MNA and succinate at 0.2 and 1 mM. Succinate was used as enhancer positive control, and 1-MNA as negative control. It is worth noting that 1-MNA does not appear in any of our learned networks, for alignment, no-alignment, Skeleton, Augmented, or any number of parents tested. D-Xylose, and succinate at 0.2 mM appears to have an enhancer effect in the stationary phase (Supplemental File 7: Figure S7(b)), but they are not statistically significant at this concentration (Supplemental File 8: Table S1). No effect was observed on *P. aeruginosa* growth in the presence 1-MNA. Though at 1 mM concentration, D-Xylose, and succinate produce an enhance the growth and it is statistically significant (Table 1) (t-test, p<0.05). The presence of 1-MNA did not have a significant effect on *P. aeruginosa* growth, it could potentially be an inhibitory compound (Figure 6).

## Data availability

All code, networks, and longitudinal microbiome data sets can be downloaded from GitHub upon publication.

## Data citation

All data analyzed in this work are derived from the iHMP IBD website: https://www.ibdmdb.org (13).

## SUPPLEMENTAL MATERIAL

**FIG S1.** Supplemental file 1 shows an adjacency matrix representation between microbiome entities for the two multi-omic frameworks used in this study: Skeleton (a) and Augmented (b). Figure highlights in red the added interactions in Augmented framework when compared to Skeleton framework.

**FIG S2.** Supplemental file 2 shows a two-stage DBN learned on the IBD data set by PALM with Skeleton constraints and a maximum number of parents of 3. Nodes are either taxa (circles), genes (diamonds), metabolites (squares), host genes (hexagons), and environmental variables (triangles). The different node types have been grouped in different circles, their transparency is proportional to their average abundance relative to that node type. While there are two consecutive time slices *t_i_* (blue) and *t_i_* _+1_ (orange), nodes with no neighbors and self loops were removed for simplicity. Dotted lines denote *intra edges* (i.e., directed links between nodes in same time slice), whereas solid lines denote *inter edges* (i.e., directed links between nodes in different time slices). Edge color indicates positive (green) or negative (red) temporal influence, and edge transparency indicates strength of bootstrap support. Edge thickness indicates statistical influence of regression coefficient after normalizing for parent values, as described in (26).

**FIG S3.** Supplemental file 3 shows a two-stage DBN learned on the IBD data set by PALM with Augmented constraints and a maximum number of parents of 3. Nodes are either host genes (hexagons), taxa (circles), genes (diamonds), or metabolites (squares). The different node types have been grouped in different circles, their transparency is proportional to their average abundance relative to that node type, and the two time slices were separated. Dotted lines denote *intra edges*, whereas solid lines denote *inter edges*. Edge color indicates positive (green) or negative (red) temporal influence and edge transparency indicates strength of bootstrap support. Edge thickness indicates statistical influence of regression coefficient after normalizing for parent values, as described in (26).

**FIG S4.** Supplemental file 4 shows the observed and predicted microbial composition trajectories for a representative aligned subject (C3028). Microbiota composition profile for this subject is comprised of the top 15 most abundant bacteria along with all remaining bacteria merged into the “other” category. The *y* axis corresponds to the relative abundance of each bacteria, while the *x* axis represents the original measured time point after alignment. Figure highlights the observed and predicted trajectories of this subject between taxa-based alignment (left) and gene-based alignment (right). We note that aligned interval for gene-based alignment is stretched and shifted when compared to the taxa-based alignment. For each alignment type, a DBN was learned with the Skeleton framework and a maximum number of parents of 3, and tested on the previously unseen C3028 subject. Gene-based alignment exhibits a lower prediction error (MAE=0.0043) than taxa-based alignment (MAE=0.0054). In this example, taxa-based alignment does a worse job at predicting low abundance bacteria than gene-based alignment.

**FIG S5.** Supplemental file 5 shows the MAE of PALM models (Augmented and Skeleton) against a baseline method (Baseline) and a previously published approach (MTPLasso) for a sampling rate of two weeks which most closely resembles the originally measured time points. Although baseline method uses only metagenomic data, gene- and metabolite-based alignment were generated using gene expression and metabolite intensities data, respectively.

**FIG S6.** Supplemental file 6 shows the *In silico* validation results with 100 bootstrap repetitions. Graphs from **(a)** and **b** show the performance of different alignment types, while **(c)** and **(d)** vary the number of parents used when learning the networks. The left part of each subfigure shows the precision (percentage of predicted edges that were validated) and the right part shows the probability of validating at least that many edges by chance (*y* axis in reverse logarithm scale so higher is better for both). The *x* represents the bootstrap value threshold that was used to select the edges included in the analysis. For example, for a threshold of 0.7, the score for edges that appear in more than 70% of the repetitions is shown. The dashed lines (*T* → *G* interactions) were learned using the *Skeleton* constraints, and the solid lines (*T* → *M* interactions) were learned using the *Augmented* constraints, because the edges *T* → *M* are not allowed directly in *Skeleton*. **(a)** Validation for *T* → *G* interactions (bacterial taxon expressing a gene) varying the alignment reference used. Noalignment barely does better than the random baseline, followed closely by the metabolite-based alignment. Taxon-based alignment has a slight better precision than gene-based, but the latter has a much better probability score than the former. **(b)** Validation for *T* → *M* interactions (bacterial taxon consuming a metabolite) varying the alignment reference used. Taxon- and metabolite-based alignment have a lower precision than the random baseline, but a better probability score. **(c)** Validation for *T* → *G* interactions (bacterial taxon expressing a gene) varying the maximum number of parents allowed. Learning with 3 Parents has a much better precision than with 4 and 5, and a similar probability score. **(d)** Validation for *T* → *M* interactions (bacterial taxon consuming a metabolite) varying the maximum number of parents allowed. Learning with 3 Parents has a better precision than with 4 and 5 for small and big thresholds. Learning with 5 parents has a better probability score for low thresholds, but it seems that it is by chance, because as the thresholds becomes more stringent, it quickly fares worse, while 3 parents overtakes 4 parents by a small percentage.

**FIG S7.** Supplemental file 7 shows the growth curves (0.2mM) of selected predicted interactions. The exponential phase was repeated and averaged over 35 replicas (up to time 14h), and after that 10 replicas were used after adding each of the metabolites at a concentration of 0.2 mM of metabolites. **(a)** *E. coli*, with Glucose and D-Xylose as positive controls **(b)** *P. aeruginosa*, wich Succinate as positive control and 1-MNA as negative control.

**TABLE S1.** Supplemental file 8 shows the metabolite effect at 0.2 mM on the two species tested. Taxa density appears in black, while the p-values are inside parenthesis. Red p-values represent a significant difference compare to LB (p<0.05). Green p-values represent a non-significant difference from LB.

**FIG S8.** Supplemental file 9 shows that the edge *Streptococcus parasanguinis* → *Pseudomonas unclassified* on the bottom gets explained when added mult-iomic data (TT stands for Taxon-Taxon network). In the multi-omic network that interaction gets replaced by *Streptococcus parasanguinis* (T) → rna polymerase (G) → D-Xylose (M) → *Pseudomonas unclassified* (T).

**FIG S9.** Supplemental file 10 shows the execution time for different maximum number of parents. The figure shows experimentally that the execution time grows linearly with the number of parents. Note that while the times shown are for 1 repetition, the DBN figures shown are with 100.

**FIG S10.** Supplemental file 11 shows the average predictive accuracy of the learned DBNs with PALM under the *Augmented* framework as a function of the maximum number parents. For each parent choice, we show the MAE using the learned DBNs from unaligned and aligned data. Observed that temporal alignments reduces MAE across all parent configurations. Additionally, the learned network for 3 parents shows the lowest average error (0.00427) over the others choices: 0.00455 for 4 parents and 0.00473 for 5 parents. It is worth highlighting that larger values for maximum number of parents does not show a MAE improvement while significantly increasing runtime for learning the DBN structure as shown in Figure S9.

**FIG S11.** Supplemental file 12 shows the average predictive accuracy of the learned DBNs with PALM under the *Skeleton* framework as a function of the maximum number parents. For each parent choice, we show the MAE using the learned DBNs from unaligned and aligned data. Observed that temporal alignments reduces MAE across all parent configurations. Additionally, the learned network for 3 parents shows the lowest average error (0.00400) over the others choices: 0.00409 for 4 parents and 0.00414 for 5 parents. It is worth highlighting that larger values for maximum number of parents does not show a MAE improvement while significantly increasing runtime for learning the DBN structure as shown in Figure S9.

**FIG S12.** Supplemental file 13 shows the *In silico* validation results of a sampling rate of 14 days compared with one day. The left part of each subfigure shows the precision (percentage of predicted edges that were validated) and the right part shows the probability of validating at least that many edges by chance (*y* axis in reverse logarithm scale so higher is better for both). The *x* represents the bootstrap value threshold that was used to select the edges included in the analysis. Green represents the original sampling rate of 14 days, while gray the option of using a sampling rate of one day. **(a)** Validation for *T* → *G* interactions (bacterial taxon expressing a gene). Both the precision and probability are very similar for both sampling rates, 14d outperforming 1d by a small amount. **(b)** Validation for *T* → *M* interactions (bacterial taxon consuming a metabolite). The 14d sampling rate option clearly outperforms the smaller sampling rate dataset, with both better precision and probability.

**FIG S13.** Supplemental file 14 shows the validation results of our normalization compared with the LogRatio normalization. The left part of each subfigure shows the precision (percentage of predicted edges that were validated) and the right part shows the probability of validating at least that many edges by chance (*y* axis in reverse logarithm scale so higher is better for both). The *x* represents the bootstrap value threshold that was used to select the edges included in the analysis. Green and blue represent our normalization for alignment and no alignment respectively, while yellow and red represent the LogRatio normalization, for alignment and no alignment respectively. **(a)** Validation for *T* → *G* interactions (bacterial taxon expressing a gene). The two normalization metrics are very similar to each other in term of precision, but when the more meaningful metric of the statistical probability tears them appart, with our normalization having a more meaningful probability by a small margin. **(b)** Validation for *T* → *M* interactions (bacterial taxon consuming a metabolite). As expected, The alignment version does better than no-alignment, for both normalization methods. Our normalization also has a better precision and statistical significance than the LogRatio normalization,

**FIG S14.** Supplemental file 15 shows the RMSE comparison between MMvec and PALM when predicting metabolites. Gene-aligned comparison between MMvec and PALM with Augmented and Skeleton restrictions for the whole dataset (All). In addition, we also compare MMvec with our learned DBNs with the Augmented constraints restricted to the dataset with just taxa and metabolites (TM) in order to make the comparisons more fair. PALM greatly outperforms MMvec at the task at hand (p-value = 4.10E-30 for a two-tailed paired t-test against TM_Augmented) for every version tried. MMvec was executed with 100.000 epochs, five latent dimensions, a learning rate of 0.00001, and a batch size of 500, leaving all other parameters to default. After executing it five times, the execution with lowest RMSE was chosen and compared against our method. The CV RMSE of the last 51 epochs (same as the number of subjects) are represented, so because by this time the algorithm had already converged, the error variance of MMvec seems small

## ACKNOWLEDGMENTS

This work was partially supported by McDonnell Foundation (ZB-J), NSF DBI-1356505 (ZB-J), NIH 1R15AI128714-01 (GN), and the FIU Dissertation Year Fellowship (DR-P). The funders had no role in study design, data collection and interpretation, or the decision to submit the work for publication.

GN and ZB-J conceived the experiments. DR-P, JL-M, GN and ZB-J designed the experiments. DR-P and JL-M performed the experiments. DR-P, JL-M, GN and ZB-J analyzed the data. DR-P performed the computational validation. NB and KM performed the biological validation. DR-P, JL-M, GN, ZB-J, and NB contributed to writing the manuscript. All authors read and approved the final manuscript.

The authors thank Shekhar Bhansali and Maximiliano S. Perez for their support.

