## Supplemental material for "Dynamic Bayesian networks for integrating multi-omics time-series microbiome data"

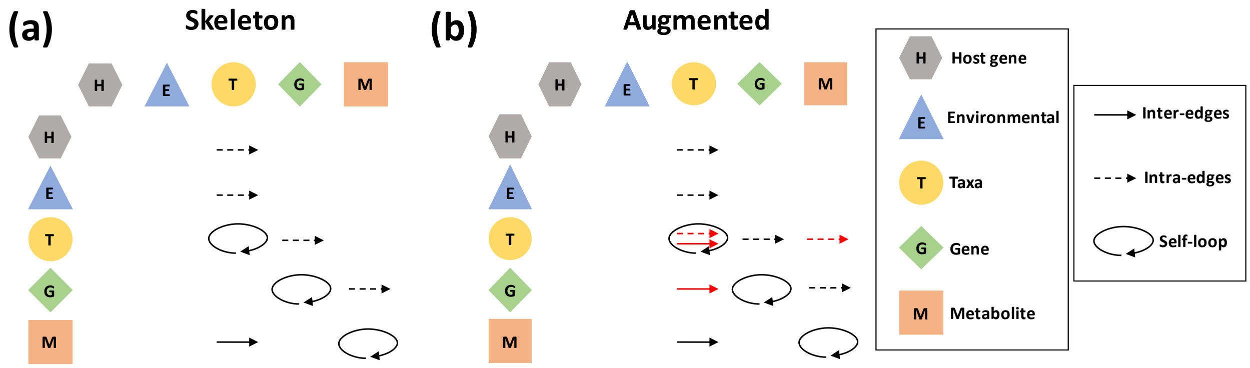


Figure S1: **Multi-omic frameworks used in this study.** Figure shows an adjacency matrix representation between microbiome entities for the two multi-omic frameworks used in this study: *Skeleton* (a) and *Augmented* (b). Figure highlights in red the added edge types in Augmented framework when compared to Skeleton framework.

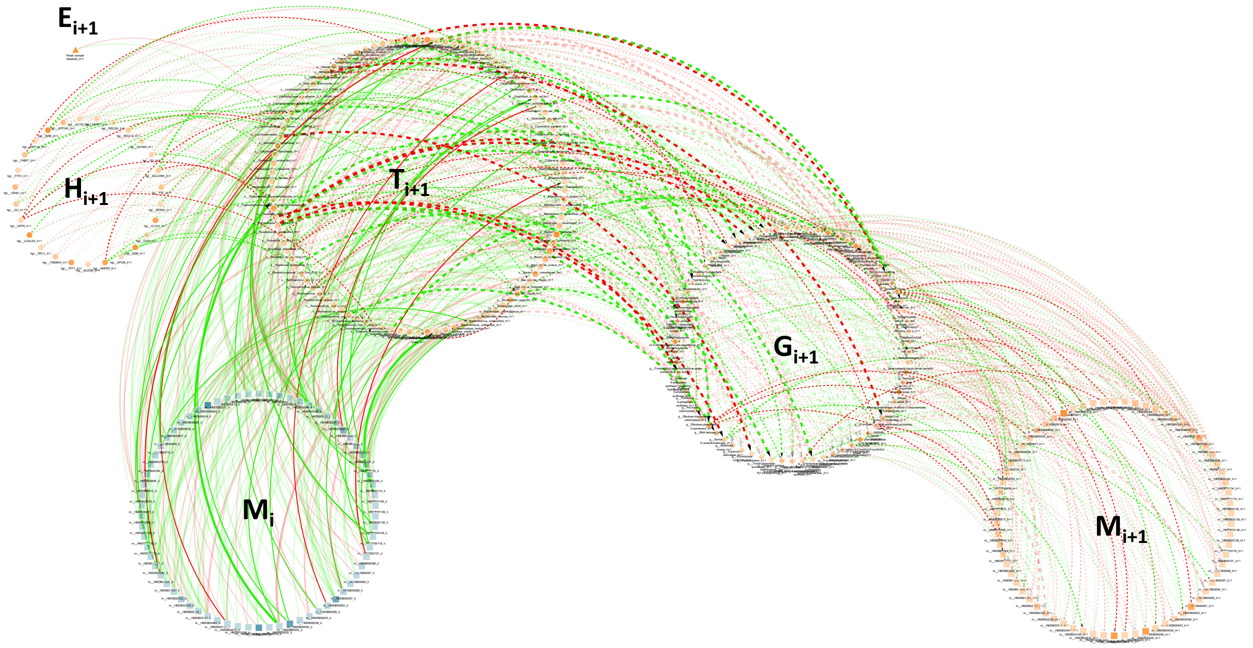


Figure S2: **Learned DBN by PALM with the Skeleton framework on the IBD data set.** Figure shows a two-stage DBN learned by PALM with Skeleton constraints and a maximum number of parents of 3. Nodes are either taxa (circles), genes (diamonds), metabolites (squares), host genes (hexagons), and environmental variables (triangles). The different node types have been grouped in different circles, their transparency is proportional to their average abundance relative to that node type. While there are two consecutive time slices $t_i$ (blue) and $t_{i+1}$ (orange), nodes with no neighbors and self loops were removed for simplicity. Dotted lines denote *intra edges* (i.e., directed links between nodes in same time slice), whereas solid lines denote *inter edges* (i.e., directed links between nodes in different time slices). Edge color indicates positive (green) or negative (red) temporal influence, and edge transparency indicates strength of bootstrap support. Edge thickness indicates statistical influence of regression coefficient after normalizing for parent values.

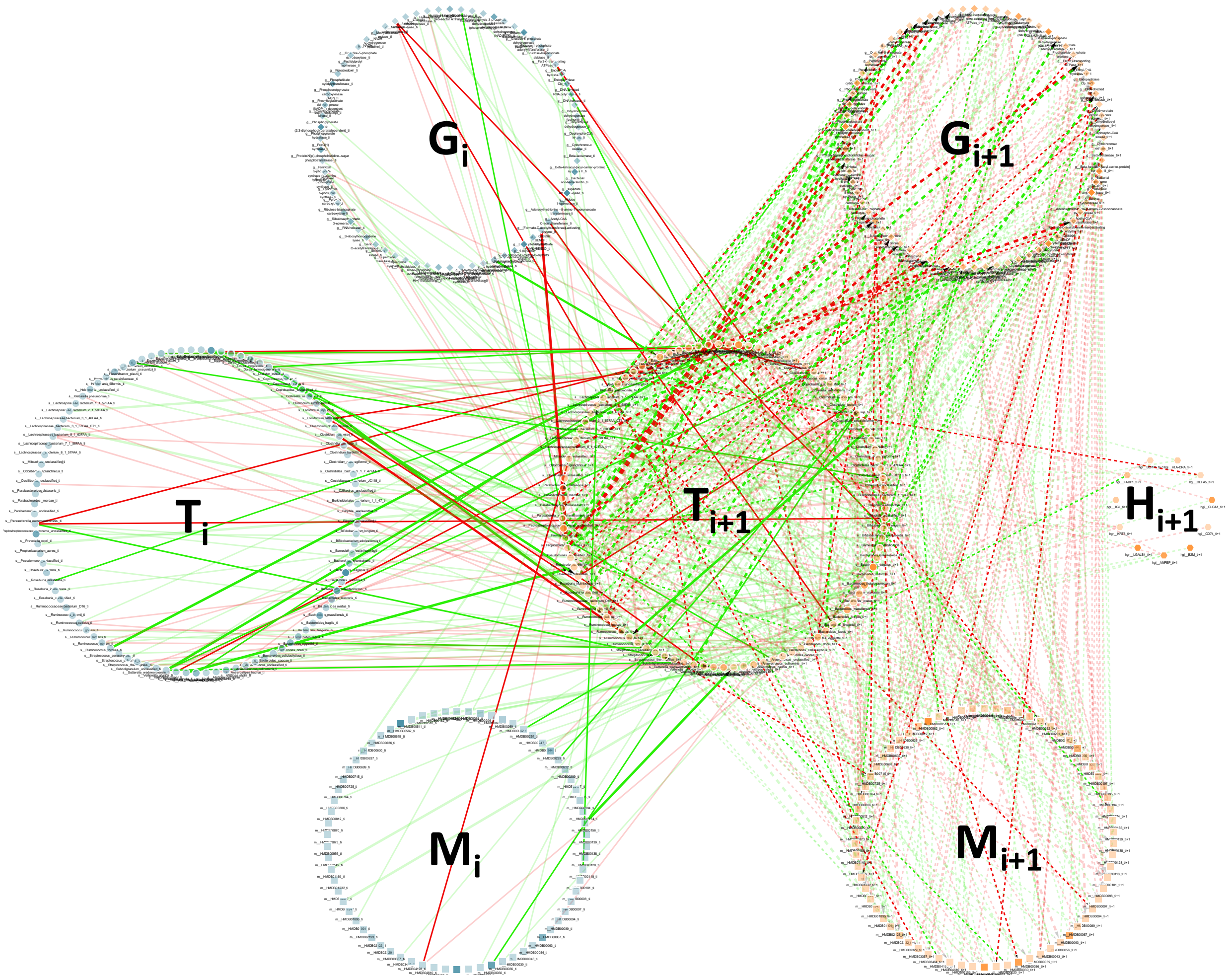


Figure S3: **Learned DBN by PALM with the Augmented framework on the IBD data set.** Figure shows a two-stage DBN learned by PALM with Augmented constraints and a maximum number of parents of 3. Nodes are either host genes (hexagons), taxa (circles), genes (diamonds), or metabolites (squares). The different node types have been grouped in different circles, their transparency is proportional to their average abundance relative to that node type, and the two time slices were separated. Dotted lines denote *intra edges*, whereas solid lines denote *inter edges*. Edge color indicates positive (green) or negative (red) temporal influence and edge transparency indicates strength of bootstrap support. Edge thickness indicates statistical influence of regression coefficient after normalizing for parent values.

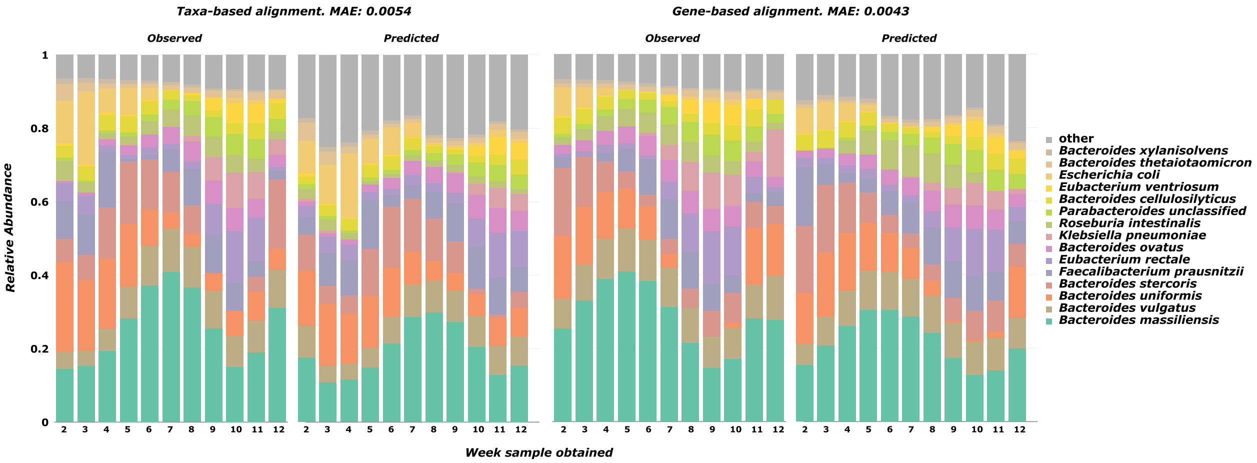


Figure S4: **Comparison of observed versus predicted microbial composition trajectories.** Figure shows the observed and predicted microbial composition trajectories for a representative aligned subject (C3028). Microbiota composition profile for this subject is comprised of the top 15 most abundant bacteria along with all remaining bacteria merged into the "other" category. The $y$ axis corresponds to the relative abundance of each bacteria, while the $x$ axis represents the original measured time point after alignment. Figure highlights the observed and predicted trajectories of this subject between taxa-based alignment (left) and gene-based alignment (right). We note that aligned interval for gene-based alignment is stretched and shifted when compared to the taxa-based alignment. For each alignment type, a DBN was learned with the Skeleton framework and a maximum number of parents of 3, and tested on the previously unseen C3028 subject. Gene-based alignment exhibits a lower prediction error (MAE=0.0043) than taxa-based alignment (MAE=0.0054). In this example, taxa-based alignment does a worse job at predicting low abundance bacteria than gene-based alignment.

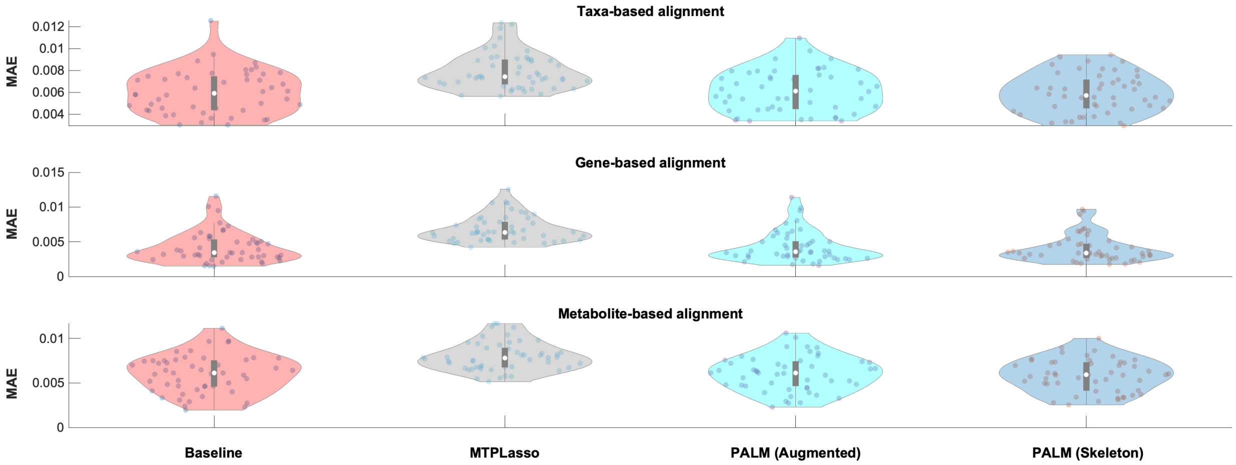


Figure S5: **Comparison of average predictive accuracy between methods on the IBD data sets aligned using taxa, gene and metabolite data.** Figure shows the MAE of PALM models (Augmented and Skeleton) against a baseline method and a previously published approach (MTPLasso) for a sampling rate of two weeks which most closely resembles the originally measured time points. Although baseline method uses only metagenomic data, gene- and metabolite-based alignment were generated using gene expression and metabolite intensities data, respectively.

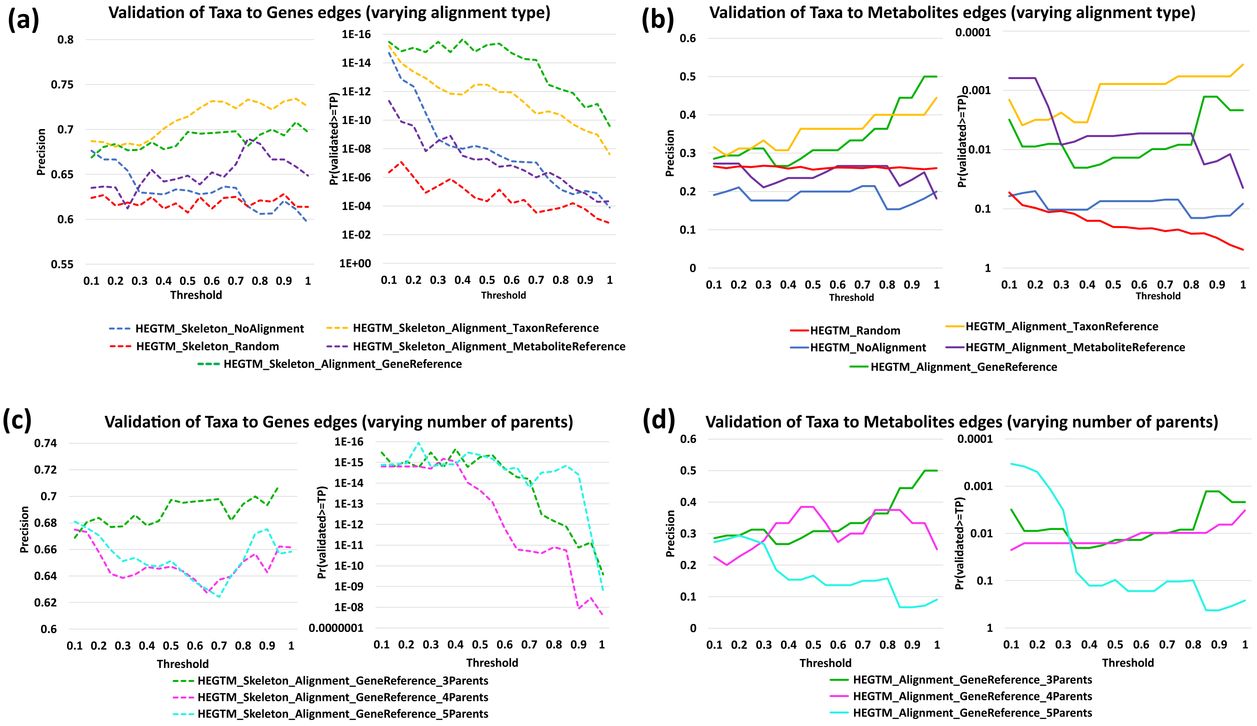


Figure S6: *In silico* **validation results with 100 bootstrap repetitions**. Graphs from **(a)** and **b** show the performance of different alignment types, while **(c)** and **(d)** vary the number of parents used when learning the networks. The left part of each subfigure shows the precision (percentage of predicted edges that were validated) and the right part shows the probability of validating at least that many edges by chance ($y$ axis in reverse logarithm scale so higher is better for both). The $x$ represents the bootstrap value threshold that was used to select the edges included in the analysis. For example, for a threshold of 0.7, the score for edges that appear in more than 70% of the repetitions is shown. The dashed lines ($T \rightarrow G$ interactions) were learned using the *Skeleton* constraints, and the solid lines ($T \rightarrow M$ interactions) were learned using the *Augmented* constraints, because the edges $T \rightarrow M$ are not allowed directly in *Skeleton*. **(a)** Validation for $T \rightarrow G$ interactions (bacterial taxon expressing a gene) varying the alignment reference used. Noalignment barely does better than the random baseline, followed closely by the metabolite-based alignment. Taxon-based alignment has a slight better precision than gene-based, but the latter has a much better probability score than the former. **(b)** Validation for $T \rightarrow M$ interactions (bacterial taxon consuming a metabolite) varying the alignment reference used. Taxon- and metabolite- based alignment have a lower precision than the random baseline, but a better probability score. **(c)** Validation for $T \rightarrow G$ interactions (bacterial taxon expressing a gene) varying the maximum number of parents allowed. Learning with 3 Parents has a much better precision than with 4 and 5, and a similar probability score. **(d)** Validation for $T \rightarrow M$ interactions (bacterial taxon consuming a metabolite) varying the maximum number of parents allowed. Learning with 3 Parents has a better precision than with 4 and 5 for small and big thresholds. Learning with 5 parents has a better probability score for low thresholds, but it seems that it is by chance, because as the thresholds becomes more stringent, it quickly fares worse, while 3 parents overtakes 4 parents by a small percentage.

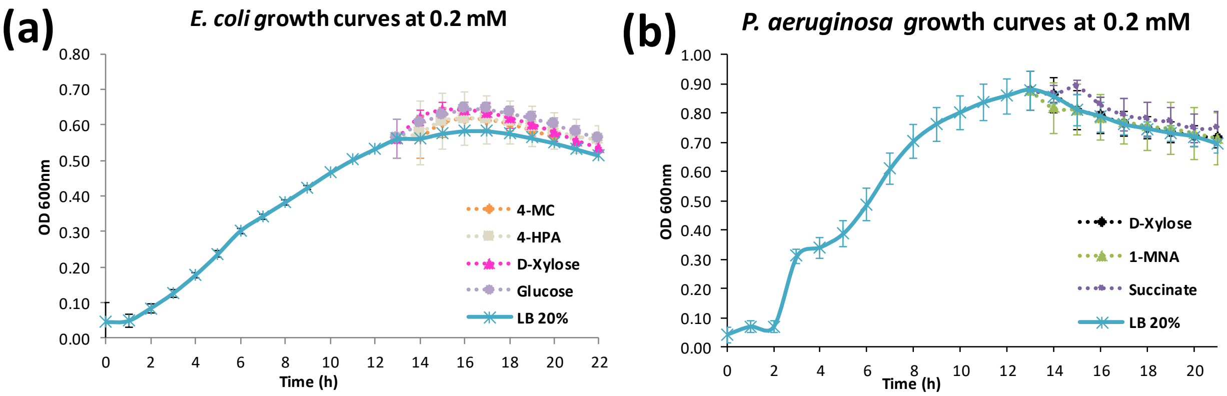


Figure S7: **Growth curves (0.2 mM)**. In this figure different metabolites were introduced at 0.2 mM concentration at the end of the exponential phase (up to time 14h). Figure shows the growth curves after all data points were averaged over 10 replicates. **(a)** *E. coli*, with Glucose and D-Xylose as positive controls. **(b)** *P. aeruginosa*, wich Succinate as positive control and 1-MNA as negative control.

| *Escherichia coli* HB101 | | | |
|---|---|---|---|
| **Met.** | **16h** | **17h** | **18h** |
| LB | 0.583 | 0.584 | 0.576 |
| 4-MC | 0.617 (0.0108) | 0.614 (0.0055) | 0.602 (0.0026) |
| 4-HPA | 0.617 (0.1458) | 0.616 (0.1332) | 0.608 (0.1049) |
| D-Xylose | 0.642 ( 0.0028) | 0.635 (0.0073) | 0.619 (0.0132) |
| Glucose | 0.644 (0.0032) | 0.644 (0.0036) | 0.633 (0.0049) |
| ***Pseudomonas aeruginosa* PAO1** | | | |
| **Met.** | **15h** | **16h** | **17h** |
| LB | 0.811 | 0.787 | 0.760 |
| D-Xylose | 0.812 (0.9029) | 0.793 (0.7561) | 0.763 (0.7710) |
| 1-MNA | 0.813 (0.9117) | 0.786 (0.9416) | 0.772 (0.6110) |
| Succinate | 0.889 (0.0128) | 0.830 (0.6979) | 0.795 (0.0530) |

Table S1: **Metabolite effect at 0.2 mM**. Taxa density appears in black, while the p-values are inside parenthesis. Red p-values represent a significant difference compare to LB ($p<0.05$). Green p-values represent a non-significant difference from LB.

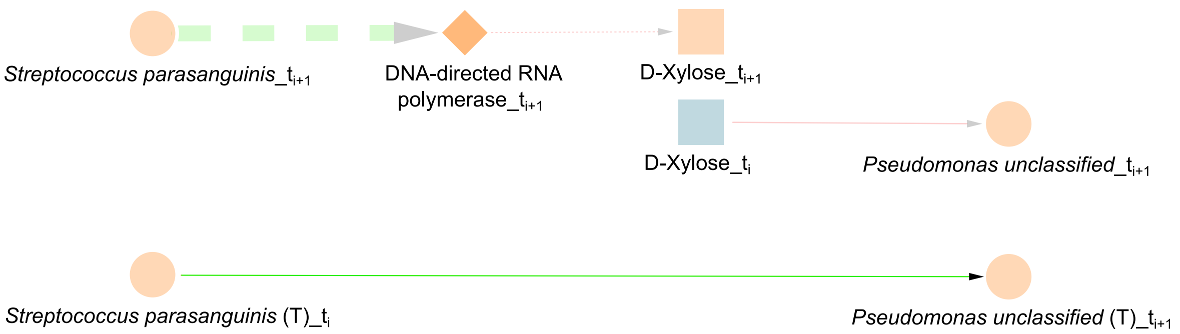


Figure S8: The edge *Streptococcus parasanguinis* → *Pseudomonas unclassified* on the bottom gets explained when added multiomic data (T stands for a dataset with just taxa). In the multi-omic network that interaction gets replaced by *Streptococcus parasanguinis* (T) → RNA polymerase (G) → D-Xylose (M) → *Pseudomonas unclassified* (T).

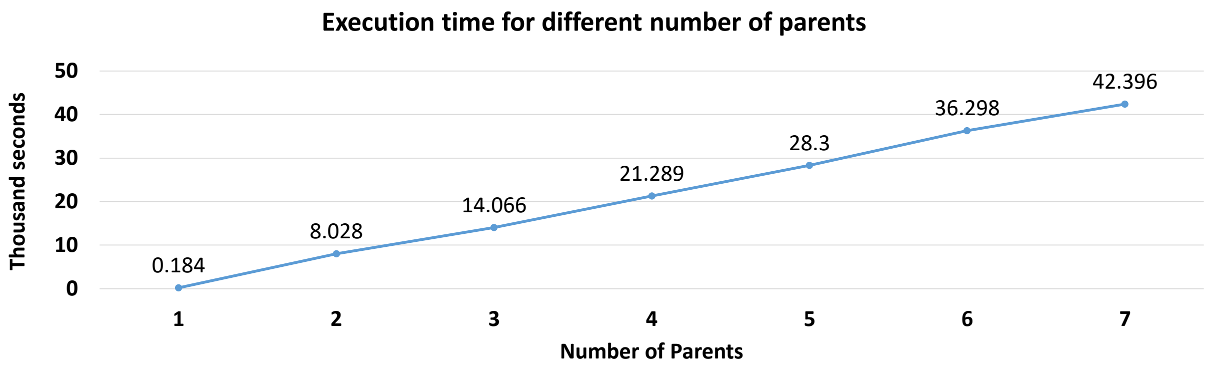


Figure S9: **Execution time for different maximum number of parents**. The figure shows experimentally that the execution time grows linearly with the number of parents. Note that while the times shown are for 1 repetition, the DBN figures shown are with 100.

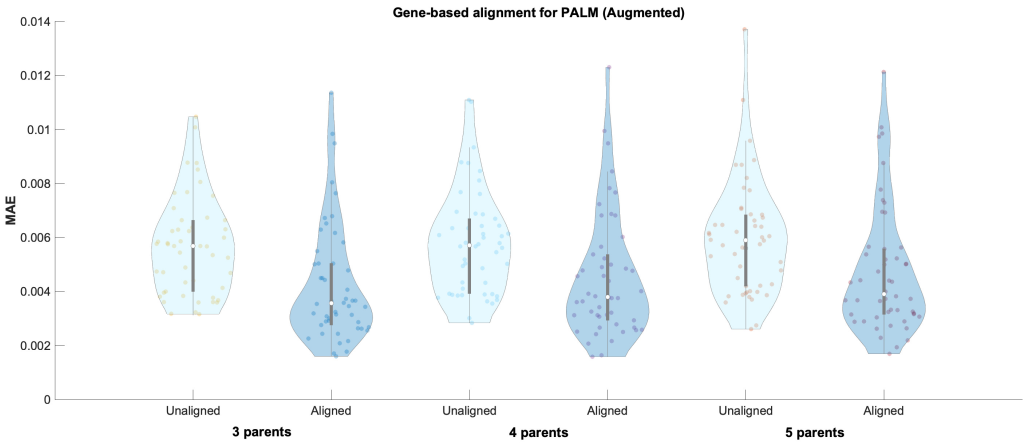


Figure S10: **MAE results a function of number of parents for the *Augmented* framework.** Figure shows the average predictive accuracy of the learned DBNs with PALM under the *Augmented* framework as a function of the maximum number parents. For each parent choice, we show the MAE using the learned DBNs from unaligned and aligned data. Observed that temporal alignments reduces MAE across all parent configurations. Additionally, the learned network for 3 parents shows the lowest average error (0.00427) over the others choices: 0.00455 for 4 parents and 0.00473 for 5 parents. It is worth highlighting that larger values for maximum number of parents does not show a MAE improvement while significantly increasing runtime for learning the DBN structure as shown in Figure S9.

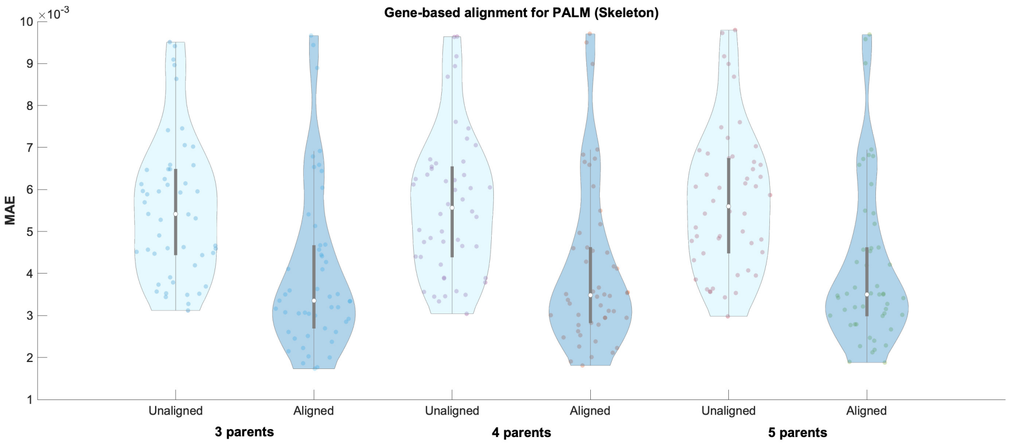


Figure S11: **MAE results a function of number of parents for the *Skeleton* framework.** Figure shows the average predictive accuracy of the learned DBNs with PALM under the *Skeleton* framework as a function of the maximum number parents. For each parent choice, we show the MAE using the learned DBNs from unaligned and aligned data. Observed that temporal alignments reduces MAE across all parent configurations. Additionally, the learned network for 3 parents shows the lowest average error (0.00400) over the others choices: 0.00409 for 4 parents and 0.00414 for 5 parents. It is worth highlighting that larger values for maximum number of parents does not show a MAE improvement while significantly increasing runtime for learning the DBN structure as shown in Figure S9.

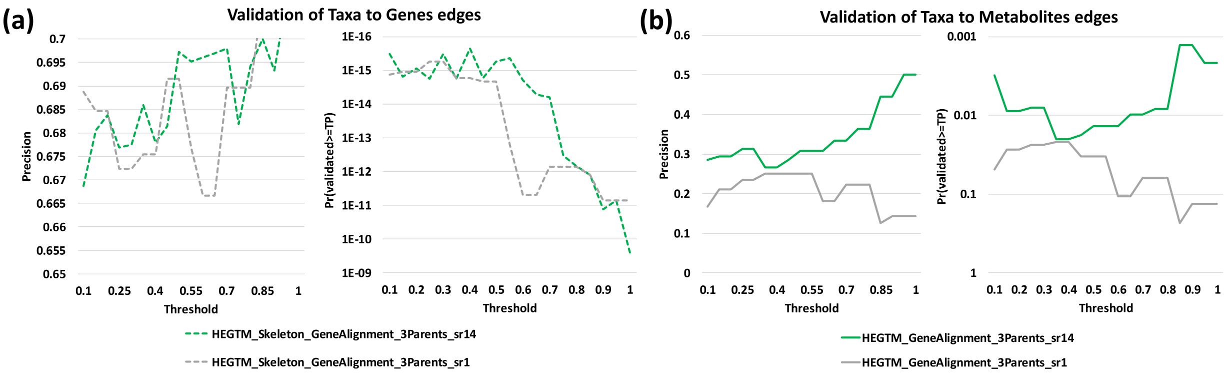


Figure S12: ***In silico* validation results of a sampling rate of 14 days compared with one day.** The left part of each subfigure shows the precision (percentage of predicted edges that were validated) and the right part shows the probability of validating at least that many edges by chance ($y$ axis in reverse logarithm scale so higher is better for both). The $x$ represents the bootstrap value threshold that was used to select the edges included in the analysis. Green represents the original sampling rate of 14 days, while gray the option of using a sampling rate of one day. **(a)** Validation for $T \rightarrow G$ interactions (bacterial taxon expressing a gene). Both the precision and probability are very similar for both sampling rates, 14d outperforming 1d by a small amount. **(b)** Validation for $T \rightarrow M$ interactions (bacterial taxon consuming a metabolite). The 14d sampling rate option clearly outperforms the smaller sampling rate dataset, with both better precision and probability.

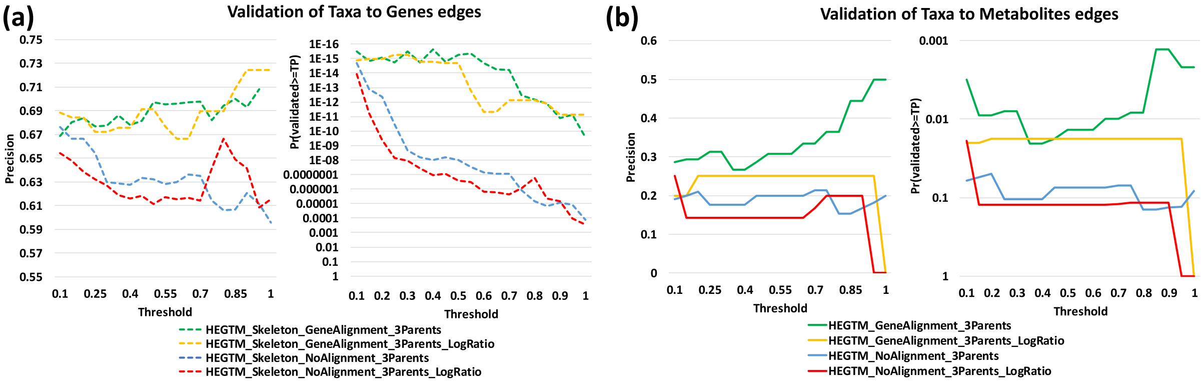


Figure S13: ***In silico* validation results of our normalization compared with the LogRatio normalization.** The left part of each subfigure shows the precision (percentage of predicted edges that were validated) and the right part shows the probability of validating at least that many edges by chance ($y$ axis in reverse logarithm scale so higher is better for both). The $x$ represents the bootstrap value threshold that was used to select the edges included in the analysis. Green and blue represent our normalization for alignment and no alignment respectively, while yellow and red represent the LogRatio normalization, for alignment and no alignment respectively. **(a)** Validation for $T \rightarrow G$ interactions (bacterial taxon expressing a gene). The two normalization metrics are very similar to each other in term of precision, but when the more meaningful metric of the statistical probability tears them appart, with our normalization having a more meaningful probability by a small margin. **(b)** Validation for $T \rightarrow M$ interactions (bacterial taxon consuming a metabolite). As expected, The alignment version does better than no-alignment, for both normalization methods. Our normalization also has a better precision and statistical significance than the LogRatio normalization, regardless of alignment type.

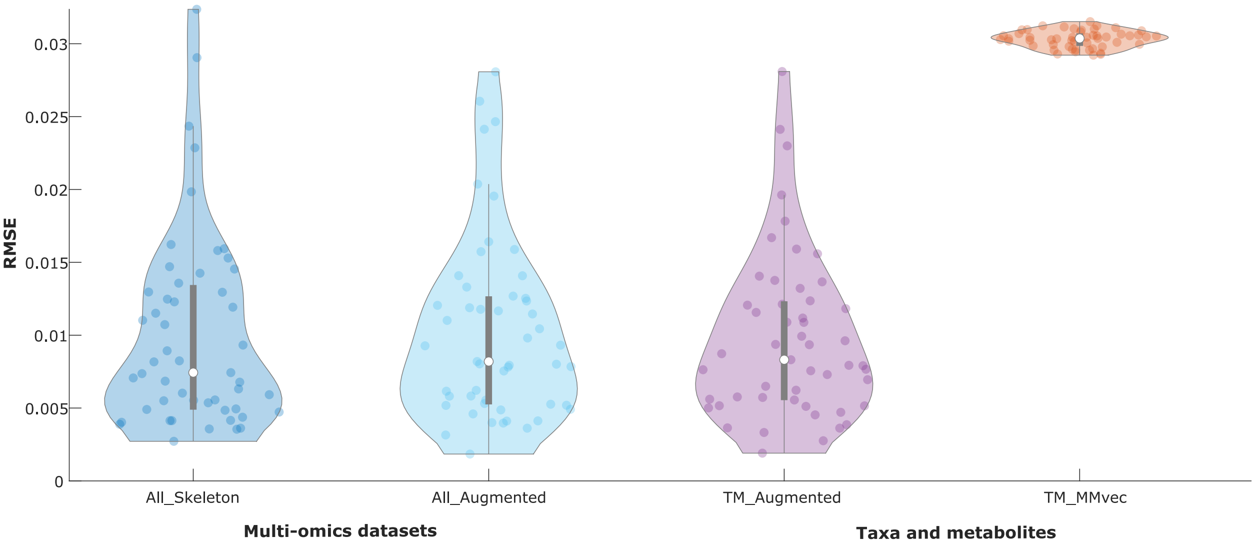


Figure S14: **RMSE comparison between MMvec and PALM when predicting metabolites**. Gene-aligned comparison between MMvec and PALM with Augmented and Skeleton restrictions for the whole dataset (All). In addition, we also compare MMvec with our learned DBNs with the Augmented constraints restricted to the dataset with just taxa and metabolites (TM) in order to make the comparisons more fair. PALM greatly outperforms MMvec at the task at hand (p-value = 4.10E-30 for a two-tailed paired t-test against TM_Augmented) for every version tried. MMvec was executed with 100.000 epochs, five latent dimensions, a learning rate of 0.00001, and a batch size of 500, leaving all other parameters to default. After executing it five times, the execution with lowest RMSE was chosen and compared against our method. The CV RMSE of the last 51 epochs (same as the number of subjects) are represented, so because by this time the algorithm had already converged, the error variance of MMvec seems small.
